# General and cell-type-specific aspects of the motor neuron maturation transcriptional program

**DOI:** 10.1101/2021.03.05.434185

**Authors:** Tulsi Patel, Jennifer Hammelman, Michael Closser, David K. Gifford, Hynek Wichterle

## Abstract

Building a nervous system is a protracted process that starts with the specification of individual neuron types and ends with the formation of mature neural circuits. The molecular mechanisms that regulate the temporal progression of maturation in individual cell types remain poorly understood. In this work, we have mapped the gene expression and chromatin accessibility changes in mouse spinal motor neurons throughout their lifetimes. We found that both motor neuron gene expression and putative regulatory elements are dynamic during the first three weeks of postnatal life, when motor circuits are maturing. Genes that are up-regulated during this time contribute to adult motor neuron diversity and function. Almost all of the chromatin regions that gain accessibility during maturation are motor neuron specific, yet a majority of the transcription factor binding motifs enriched in these regions are shared with other mature neurons. Collectively, these findings suggest that a core transcriptional program operates in a context-dependent manner to access cell-type-specific cis-regulatory systems associated with maturation genes. Discovery of general principles governing neuronal maturation might inform methods for transcriptional reprogramming of neuronal age and for improved modelling of age-related neurodegenerative diseases.

## Introduction

The development of a functional nervous system requires the specification and maturation of diverse neuron types. Most neurons are generated in a brief time window during fetal development but the nervous system continues to develop for weeks in mice and years in humans before functional outputs of the nervous system, such as movement and behavior, become adult-like (Altman and Sudarshan, 1975; Bishop, 1982a, b). During this time, young neurons undergo a complex process of migration, neurite outgrowth, synapse formation, electrophysiological maturation, neurite pruning, and modulation of synaptic strength. Although this protracted process is fundamentally important for the integration of nascent neurons into functioning neural circuits, how it is orchestrated at the molecular level remains poorly understood.

Motor neurons in the brain stem and spinal cord control all skeletal muscle movements, including walking, breathing, and fine motor skills, by communicating central motor commands to diverse muscle targets. These neurons are among the first nerve cells born during embryonic development, but motor circuits and motor behaviors continue to mature over a prolonged period, extending into the postnatal life (Altman and Sudarshan, 1975; Jessell, 2000; Smith et al., 2017). Decades of developmental studies have identified key signaling molecules and cell intrinsic transcriptional programs that specify motor neuron identity during embryonic development. These studies culminated in the discovery of motor neuron selector transcription factors, Isl1 and Lhx3, which are sufficient to ectopically activate a motor neuron gene expression program in other neural progenitors (Lee and Pfaff, 2003; Pfaff et al., 1996) and even accelerate post-mitotic specification by directly reprogramming pluripotent stem cells into motor neurons (Mazzoni et al., 2013). While some of the core motor neuron effector genes, including those that control cholinergic neurotransmission, become induced soon after progenitor cells become post-mitotic, other genes required for physiological maturation and connectivity exhibit dynamic regulation or induction only at postnatal developmental stages (Arber et al., 2000; De Marco Garcia and Jessell, 2008; Kaplan et al., 2014; Mendelsohn et al., 2017; Morisaki et al., 2016; Price et al., 2002). Understanding the signaling and transcriptional programs that control dynamic gene expression in postmitotic motor neurons is of fundamental importance. Maturation status influences not only normal motor neuron function and connectivity, but also susceptibility to degeneration. During a brief window of embryonic programmed cell death, motor neurons survival is dependent on neurotrophic signals; at perinatal stages, motor neurons show selective sensitivity to decreased levels of SMN protein in mouse models of spinal muscular atrophy; and in adult animals carrying ALS causing mutations, there is a wide range of temporal onsets of motor neuron degeneration (Kanning et al., 2010). Thus, the maturation state of motor neurons is an important variable in understanding not only the normal function of motor circuits, but also the susceptibility to cell death and disease.

To elucidate mechanisms that control maturation, we performed a detailed characterization of gene expression and chromatin accessibility in embryonic and postnatal motor neurons. To accomplish this, we relied on the purification of motor neuron nuclei from dissected and homogenized spinal cords, as it is not feasible to isolate whole postnatal motor neurons because of their large size and fragility. Our data demonstrate that the motor neuron gene expression program remains highly dynamic from embryonic stages through postnatal day 21, at which point motor circuits are also known to become fully mature (Donatelle, 1977; Smith et al., 2017). We identified motor neuron specific and shared aspects of neuronal maturation by comparing the maturation trajectory of motor neurons to those of cortical neurons (Stroud et al., 2020). Analysis of accessible chromatin regions, many of which function as cis-regulatory elements controlling gene expression, identified both motor neuron specific and shared neuronal regulators that likely control the dynamic transcriptional changes during maturation. This work provides a framework for studying mechanisms that govern neuronal maturation, paving a way for better understanding age-related pathologies and neurodegeneration.

## Results

### Mapping temporal gene expression profiles of mouse motor neurons

Cell-type-specific gene expression profiles are controlled by the binding of transcription factors at cis-regulatory elements. In order to elucidate mechanisms that control motor neuron maturation, we therefore assembled a detailed temporal trajectory of gene expression and chromatin accessible regions in mouse motor neurons. To do this, we isolated spinal cord motor neurons from embryonic (E10.5, E13.5), neonatal (P4), juvenile (P13, P21), adult (P56), and aged (2 year) mice for RNA-seq and ATAC-seq analysis (**Fig. 1a**). These timepoints represent key milestones in motor neuron maturation and disease: between E10.5-E13.5 nascent motor neurons lose progenitor identity, extend their axons towards their cognate muscle targets and start to express subtype specific markers, between P4-P21 local spinal interneuron circuits mature, descending motor circuits are wired, and adult-like movement is acquired (Altman and Sudarshan, 1975; Bucchia et al., 2018; Jessell, 2000). In more aggressive mouse models of ALS (SOD1^G93A^), cellular pathologies such as swollen mitochondria, ubiquitinated protein aggregates and ER stress are apparent by ~P56, with phenotypes getting progressively worse till substantial motor neuron loss and muscle paralysis is observed at ~P120 (Kanning et al., 2010).

**Figure 1:**
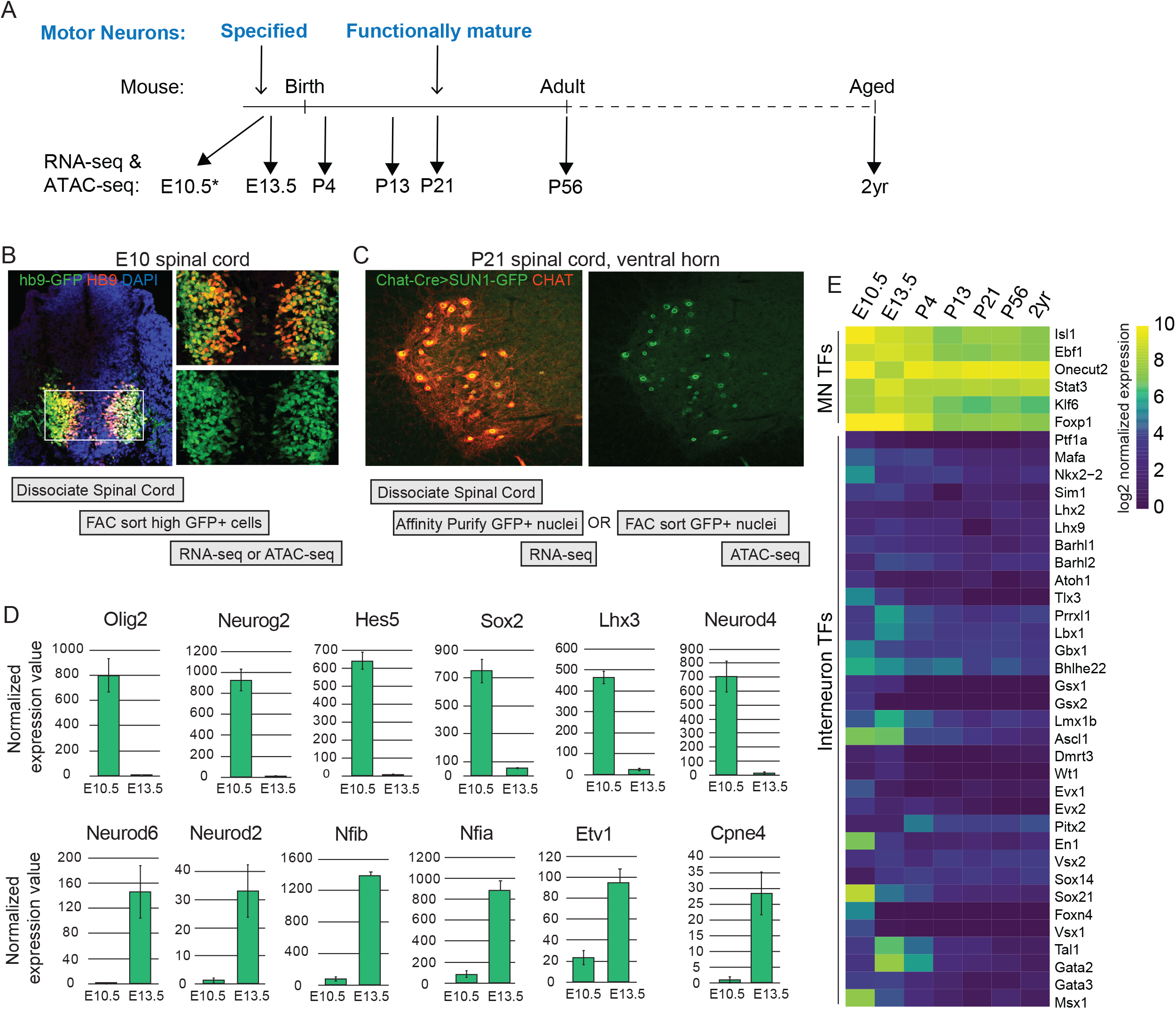
Purification of spinal motor neurons at embryonic and postnatal ages. A) Motor neurons (MNs) are specified at E10.5, become functionally mature at ~P21, and are maintained for the rest of life. We isolated whole motor neurons at E10.5(*) and motor neuron nuclei at all later timepoints to perform RNA-seq and ATAC-seq analysis. B) A spinal cord section from an Hb9-GFP E10 embryo and outline of the procedure used to isolate motor neurons. Immunostaining shows co-labeling of HB9 protein (red) with high GFP expression from the transgene (green). The boxed region from the left image is shown with split colors on the right. C) The right ventral horn region of a P21 Chat-Cre; Sun1-GFP spinal cord and outline of the procedure used to isolate motor neuron nuclei at all ages from E13.5 to 2yrs. Immunostaining shows co-labeling of CHAT protein (red) with SUN1-GFP (green). D) Normalized expression values of genes previously shown to be downregulated between E10.5 and E13.5 (top row), or upregulated between E10.5 and E13.5 (bottom row). E) Heatmap showing expression of motor neuron transcription factors and spinal interneuron transcription factors at all profiled ages.

We used two mouse lines to isolate motor neurons: the Hb9:GFP transgenic line which labels motor neuron cells starting at a nascent stage, and Chat-Cre; Sun1-sfGFP-Myc which continuously labels motor neuron nuclei at all ages after ~E12.5 (Mo et al., 2015; Rossi et al., 2011; Wichterle et al., 2002). For gene expression and chromatin accessibility analysis at E10.5, spinal cords from Hb9:GFP mice were dissociated and whole motor neuron cells expressing GFP were purified by fluorescent activated cell sorting (FACS) for RNA-seq and ATAC-seq (**Fig. 1b**, Methods). Because purification of whole motor neurons from postnatal ages is highly inefficient, we relied on the Sun1-GFP tag to isolate motor neuron nuclei from spinal cords at all ages from E13.5 to 2yrs. To do this, spinal cords were dissected from mice carrying Chat-Cre driven expression of Sun1-GFP. GFP expressing nuclei were either affinity purified using an optimized INTACT procedure for RNA-seq or FACS purified for ATAC-seq (**Fig. 1c**, Methods) (Mo et al., 2015).

To ensure that RNA-seq data collected from whole cells at E10.5 are comparable to data collected from nuclei at later ages, we examined the expression of genes that are known to be differentially expressed between E10.5 and E13.5. Previous studies have shown that nascent E10.5 motor neurons retain expression of progenitor markers such as Olig2 and Sox2, while motor neurons at E13.5 start to acquire subtype identities and expression of pool-specific markers such as Etv1 and Cpne4 (Arber et al., 2000; Delile et al., 2019; Mendelsohn et al., 2017). We examined the expression of genes known to be dynamic between E10.5 and E13.5 and observed the expected relative expression patterns in our dataset (**Fig. 1d**). This finding is in agreement with prior studies which established a high concordance between nuclear and whole cell transcriptomes (Lake et al., 2018).

To determine the overall purity of motor neurons isolated at all ages, we examined the expression of spinal motor neuron and spinal interneuron transcription factors in our dataset (Bikoff et al., 2016; Catela et al., 2019; Dasen et al., 2008; Delile et al., 2019; Ericson et al., 1992; Francius and Clotman, 2010; Hoang et al., 2018; Laub et al., 2001; Lee et al., 2013; Velasco et al., 2017). Whereas motor neuron transcription factors Isl1, Ebf1, Onecut2, Stat3, Klf6, and Foxp1 are highly and consistently expressed at all timepoints, spinal interneuron transcription factors show low or inconsistent expression (**Fig. 1e**). We therefore conclude that the gene expression and chromatin accessibility data generated in this work faithfully capture the transcriptional state of spinal motor neurons at various ages.

### Gene expression in motor neurons is dynamic during functional maturation and stabilizes during the third postnatal week

In order to understand the global dynamics of gene expression during the life of motor neurons, we performed a Principal Component Analysis (PCA) on the RNA-seq data collected at all timepoints (**Fig. 2a**). The first principal component, which explains 39.6% of the overall variance, separates the gene expression data sequentially by age, with the timepoints between E10.5 - P4 separating most prominently. Next, to identify genes that are expressed dynamically with age, we performed differential gene expression analysis between subsequent ages using the EdgeR algorithm (McCarthy et al., 2012; Robinson et al., 2010) (**Fig 2b, 2c**). These analyses show that ~40% of expressed genes are significantly up or down-regulated during the transition from E10.5 to E13.5, and from E13.5 to P4 (p < 0.001, fold change >= 2). Both the percent of genes that are differentially expressed and the extent of the differential expression (in terms of p-value and fold change) decreases as motor neurons get older, with gene expression becoming stable after P21 and remaining that way for the rest of life (**Fig 2b, 2c**). Previous studies have demonstrated that the maturation of motor circuits and motor behavior occurs during the third postnatal week, between P14-P21 (Altman and Sudarshan, 1975; Bucchia et al., 2018; Smith et al., 2017). The RNA-seq data reveals a striking concordance between the timing of transcriptional maturation and functional maturation, suggesting that transcriptional changes are required for the functional output of adult motor neurons. Indeed, when we perform a pathway enrichment analysis (Jassal et al., 2020) on the top 1000 genes that are upregulated at P21 compared to E13.5 (**Supplementary Fig. 2a**), we find that most enriched pathways include functional categories such as potassium channels, protein interactions at synapses, and muscle contraction (p<0.001). On the other hand, genes downregulated with age are enriched in common cellular pathways such as RNA and protein metabolism, translation, and mitotic pathways, in addition to neurodevelopment-specific pathways such as axon guidance (p<0.0005) (**Fig. 2d, 2e**). We have confirmed the dynamic expression patterns of several postnatally upregulated genes with in situ hybridization data available from the Allen Spinal Cord Atlas (**Supplementary Fig. 1**) (Sunkin et al., 2013).

**Figure 2:**
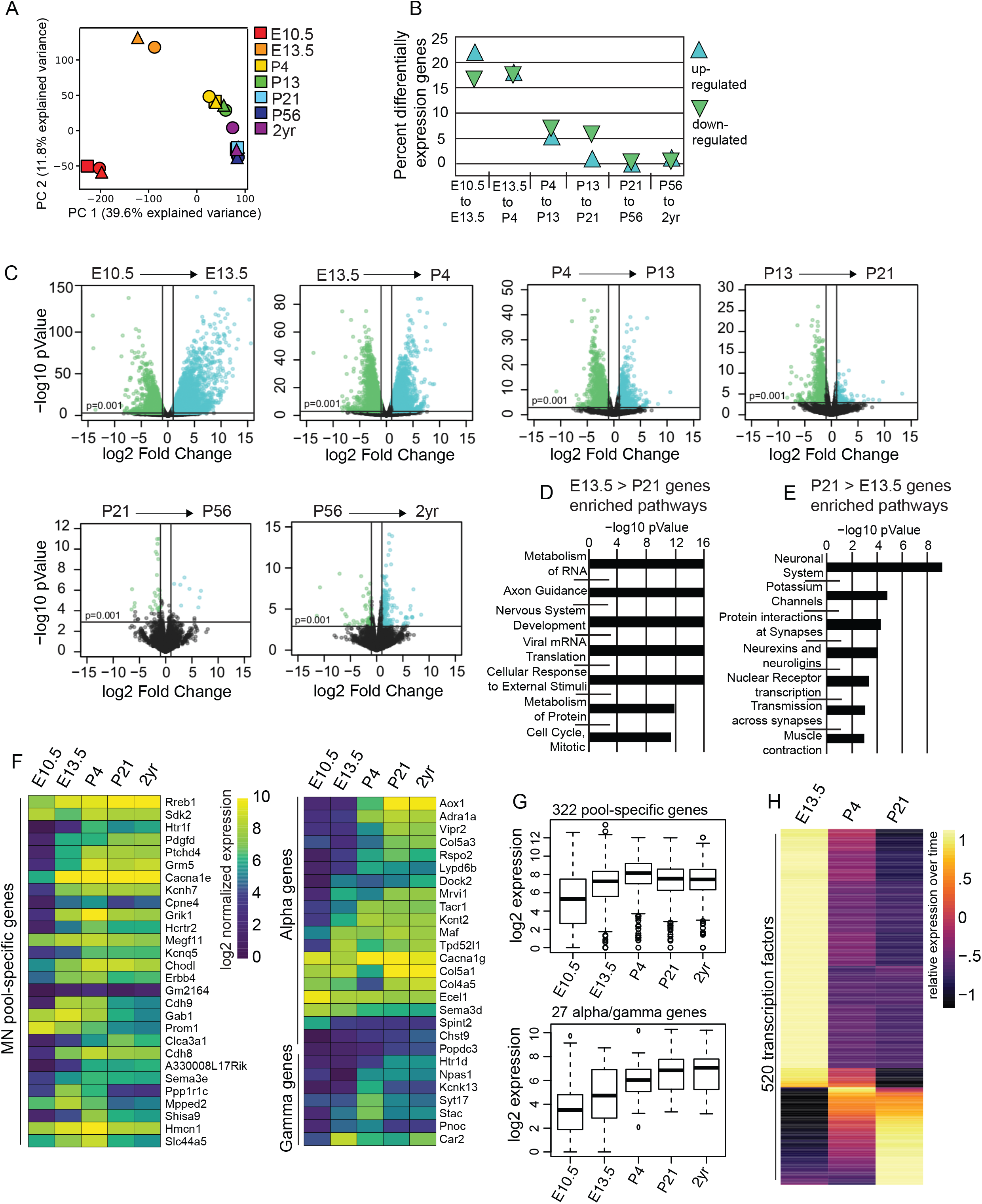
Dynamic gene expression changes during maturation contribute to adult motor neuron diversity and function. A) Principal Component Analysis on RNA-seq data at all timepoints. The plot shows the distribution of gene expression data over the first two principal components. Shapes of the same color represent biological replicates. B) Percent of expressed genes that are up-regulated or down-regulated at least 2-fold with a pValue < 0.001 (as determined by EdgeR) between two consecutive timepoints. Expressed genes have CPM >1 in at least one replicate over all timepoints. This plot is a summary of the differential gene expression plots in C. C) Plots showing differential gene expression between consecutive timepoints. Each dot represents one gene, the x-axis shows the log_2_ fold change, the y-axis shows the −log_10_ pValue. Colored dots are genes that are upregulated (blue) or downregulated (green) at least 2-fold with a pValue < 0.001 (as determined by EdgeR). D) The most significant pathways enriched in the top 1000 genes down-regulated during maturation between E13.5 and P21. E) The most significant pathways enriched in the top 1000 genes up-regulated during maturation between E13.5 and P21. F) Heatmaps showing expression of 28 motor pool specific genes (left) and 27 alpha and gamma specific genes (right) over time. Lists of markers genes were obtained from Blum et al., 2021. G) Boxplots showing expression of 322 motor pool-specific genes and 27 alpha and gamma specific genes (from F). Each plot shows the distribution of log_2_ expression values for pool markers (top plot) and alpha/gamma markers (bottom plot) over time. H) Relative expression values of 520 expressed transcription factors (Riken database) that show differential expression of at least 2-fold with a pValue < 0.001(as determined by EdgeR) between E13.5 and P21.

### Motor neuron subtype identities are temporally established during maturation

Spinal cord motor neurons are categorized into column and pool subtypes based on their rostro-caudal position and the muscle groups they innervate. Motor neurons can be further subdivided functionally into alpha, gamma, or beta types based on their electrophysiological properties and the type of muscle fiber they innervate. Developmental studies in mice and chicks have found that transcriptionally distinct column and pool identities start to be evident at E13.5 (Arber et al., 2000; De Marco Garcia and Jessell, 2008; Mendelsohn et al., 2017; Price et al., 2002). However, markers that distinguish alpha and gamma subtypes of motor neurons are not evident till late embryonic and postnatal ages (Kanning et al., 2010; Shneider et al., 2009). A recent single nucleus analysis of adult motor neurons identified several markers that distinguish motor pools and alpha vs. gamma motor neurons (Blum et al., 2021). In order to determine when subtype-specific identities are established in motor neurons, we examined the temporal trajectory of these genes in our dataset. Interestingly, we found that out of 28 pool identity markers, ~90% are upregulated between E10.5 and E13.5, 71% are induced or further upregulated between E13.5 and P4, and only 10% are upregulated between P4 and P21 (**Fig. 2f**), suggesting that motor pools identities are specified early on during embryonic and perinatal development. Analysis of the temporal expression patterns of an expanded list of 322 pool identity markers showed a similar pattern- over 90% of these genes reach their highest expression level at P4 (**Fig. 2g**). On the other hand, markers that distinguish alpha and gamma motor neurons continue to be upregulated until P21. Of 27 alpha/gamma markers, 48%, 55% and 44% are induced or upregulated between E10.5 and E13.5, E13.5 and P4, and P4 and P21, respectively (**Fig. 2f, 2g**). This difference in dynamics shows that motor pool identities, which are important for axon targeting and muscle innervation are established early while alpha vs. gamma transcriptional identities, which determine different firing properties of motor neurons, are still being refined as circuits mature in postnatal life.

### Expression of neurotransmitter pathway genes is dynamic in subsets of motor neurons

Spinal motor neurons control muscle contractions through the release of the acetylcholine neurotransmitter. Our data show that acetylcholine pathway genes are expressed continuously in motor neurons, with some changes in expression levels over time (**Supplementary Fig. 2b**). Intriguingly, we also see low levels of expression of other neurotransmitter pathway genes in maturing motor neurons. These include a transient expression of TH (tyrosine hydroxylase expressed in monoaminergic neurons) in nascent postmitotic neurons, and postnatal expression of the GABAergic genes, Gad1 and Gad2 (glutamate decarboxylase, which make GABA from glutamate), and at very low levels Slc32a1 (GABA vesicular transporter); the glycinergic gene, Slc6a5 (glycine transporter); and the glutamatergic gene, Slc17a6 (vesicular glutamate transporter, Vglut2). To confirm that these expression patterns are not a result of contamination from other cell types in our dataset, we examined the expression of these genes in the published single nucleus RNA-seq analysis of adult motor neurons (Blum et al., 2021). Interestingly, the single nucleus data demonstrate that the above genes are expressed in subsets of adult skeletal muscle neurons belonging to specific subtypes (**Supplementary Fig. 2c**). Together these data raise the possibility that small subsets of postnatal motor neurons transcribe genes that are involved in the synthesis or transport of GABA, Glycine, and Glutamate neurotransmitters. Previous studies examining the presence of glutamate transporter proteins, including Vglut2, have shown inconsistent results; however, there is evidence that motor neurons use aspartate in addition to acetylcholine for neurotransmission (Mentis et al., 2005; Nishimaru et al., 2005; Richards et al., 2014). Therefore, whether the expression patterns detected in bulk and snRNA-seq data results in neurotransmission or other metabolic functions remains to be examined.

### Changes in chromatin accessibility are correlated with dynamic gene expression

Having determined the temporal dynamics of gene expression in motor neurons, we next sought to identify the regulatory factors that orchestrate maturation. Even though similar proportions of all genes are up- and down-regulated between E13.5 and P21, we found that a majority of transcription factors are downregulated over the same time period (**Supplementary Fig. 2a, Fig. 2h**). A similar trend has been previously observed in sensory neurons as well as cortical interneurons (Sharma et al., 2020; Stroud et al., 2020), suggesting that while expression of many transcription factors is required at high levels to specify cellular identities, a smaller number of transcription factors or lower levels of expression are required for the maintenance of neuronal identity and for activation of mature gene expression programs.

To identify potential regulators of motor neuron maturation, we identified age-specific accessible chromatin regions by performing ATAC-seq on E10.5, E13.5, P4, P13, P21, P56, and 2yr old motor neurons (Buenrostro et al., 2015; Corces et al., 2017). We identified >100,000 accessible peaks at each timepoint (Methods). Principal Component Analysis of the temporal ATAC-seq data demonstrated that similar to gene expression changes, regulatory regions are dynamic between E10.5-P13, with all the remaining postnatal ages clustering together (**Fig. 3a**). Calculating the percent of differentially accessible peaks between subsequent timepoints revealed that chromatin regions are extremely dynamic at early ages, with up to 50% of total ATAC peaks behaving dynamically between E10.5 and E13.5, and become increasingly stable as motor neurons mature (**Fig. 3b**). The vast majority of differentially accessible chromatin regions (~95%) are located at least 2 kb away from transcription start sites (**Fig 3c**), suggesting that changes in gene expression over time are regulated by distal regulatory regions rather than promoter proximal ones. A similar trend is also observed in temporally dynamic regulatory regions in postnatal cortical interneurons (Stroud et al., 2020). The temporal trajectory of dynamic chromatin regions closely correlates with the trajectory of gene expression changes. We therefore asked if genes that are upregulated during maturation are also surrounded by maturation specific accessible chromatin regions. To answer this, we examined the number of e13.5 > P21 accessible peaks (regions that are accessible at e13.5, but not at P21) and the number of P21 > e13.5 accessible peaks that can be assigned to the top 250 genes upregulated between e13.5 and P21 based on proximity. We found that 125 of the 250 upregulated genes contained more P21 > e13.5 accessible peaks, while only 29 genes contained more e13.5 > P21 accessible peaks (**Fig. 3d**). This observation is consistent with the idea that adult-specific accessible regions contribute to the developmental upregulation of motor neuron genes during maturation.

**Figure 3:**
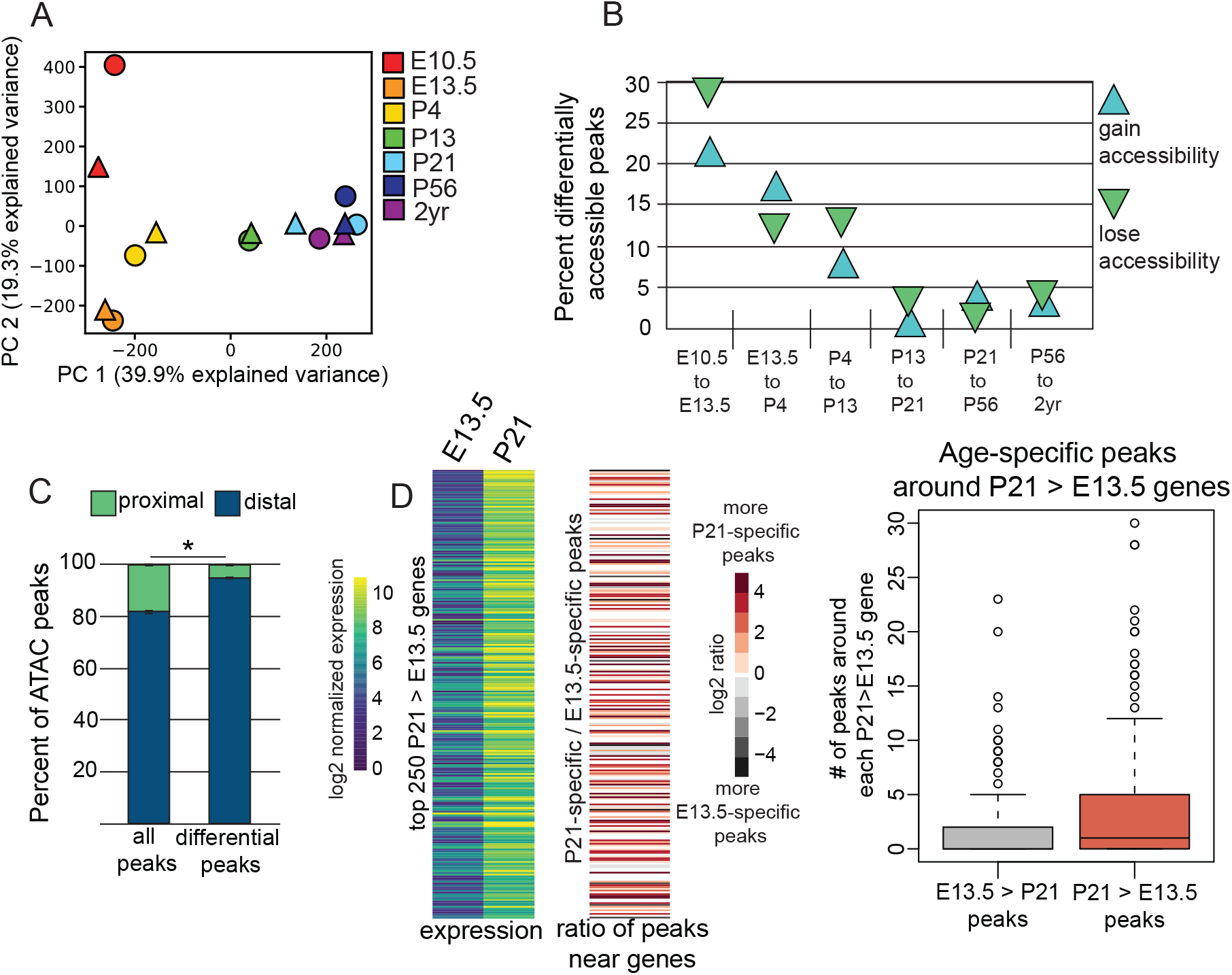
Dynamic changes in accessible regions identified by ATAC-seq correlate with dynamic gene expression. A) Principal Component Analysis on ATAC-seq data at all timepoints. The plot shows the distribution of accessibility data over the first two principal components. Shapes of the same color represent biological replicates. B) The percent of genomic regions that lose accessibility or become newly accessible between consecutive timepoints. C) The percent of total peaks or differential peaks that are proximal (within 2kb) or distal (>2kb away) from transcription start sites. For the left bar, the percent of proximal or distal peaks in the top 100k peaks at each age were calculated and averaged. For the right bar, the percent of proximal or distal peaks in ATAC-seq peaks that are differentially accessible at consecutive ages were calculated and averaged. D) The distribution of age-specific accessibility regions near genes that are upregulated during maturation. Each row across both heatmaps represents one gene. The left heatmap shows the expression at E13.5 and P21. The right heatmap shows the ratio of age specific peaks around each gene. The log_2_ ratio of peaks is calculated by dividing P21 > E13.5 peaks (peaks that are accessible at P21, but not at E13.5) by E13.5 > P21. Positive values indicate a higher number of P21 > E13.5 peaks, while negative values indicate a higher number of E13.5 > P21 peaks. The boxplot on the far right shows the number of E13.5 > P21 and P21 > E13.4 accessible regions around each of the 250 upregulated genes.

### Putative regulators of maturation can be identified from temporally accessible genomic regions

Accessible genomic regions are known to harbor binding sites for transcription factors that regulate gene expression (Buenrostro et al., 2015; Klemm et al., 2019; Mo et al., 2015; Rhee et al., 2016; Yue et al., 2014). We therefore performed de novo motif enrichment on chromatin accessible regions at each age in order to identify putative regulators of age-specific gene expression patterns. To enrich for genomic regions relevant for gene regulation during motor neuron maturation, we filtered out regions that are also accessible in embryonic stem cells (ESCs) (de Dieuleveult et al., 2016), and regions bound by CTCF, which are a hallmark of insulators (based on CTCF ChIP-seq on ESC-derived motor neurons, unpublished data), a hallmark of sites controlling genome architecture (Guo et al., 2015; Ong and Corces, 2014). Of the remaining peaks, we selected the 10k most significant accessible regions at each age to perform de novo motif enrichment analysis using HOMER (Heinz et al., 2010). These accessible genomic regions are well conserved (**Fig. 4a**), suggesting functional importance. Interestingly, we find that accessible E10.5 and E13.5 regions are the most conserved while postnatal regions are sequentially less conserved, consistent with the reported higher conservation of forebrain regulatory enhancers active at embryonic ages compared to the postnatal ones (Nord et al., 2013).

**Figure 4:**
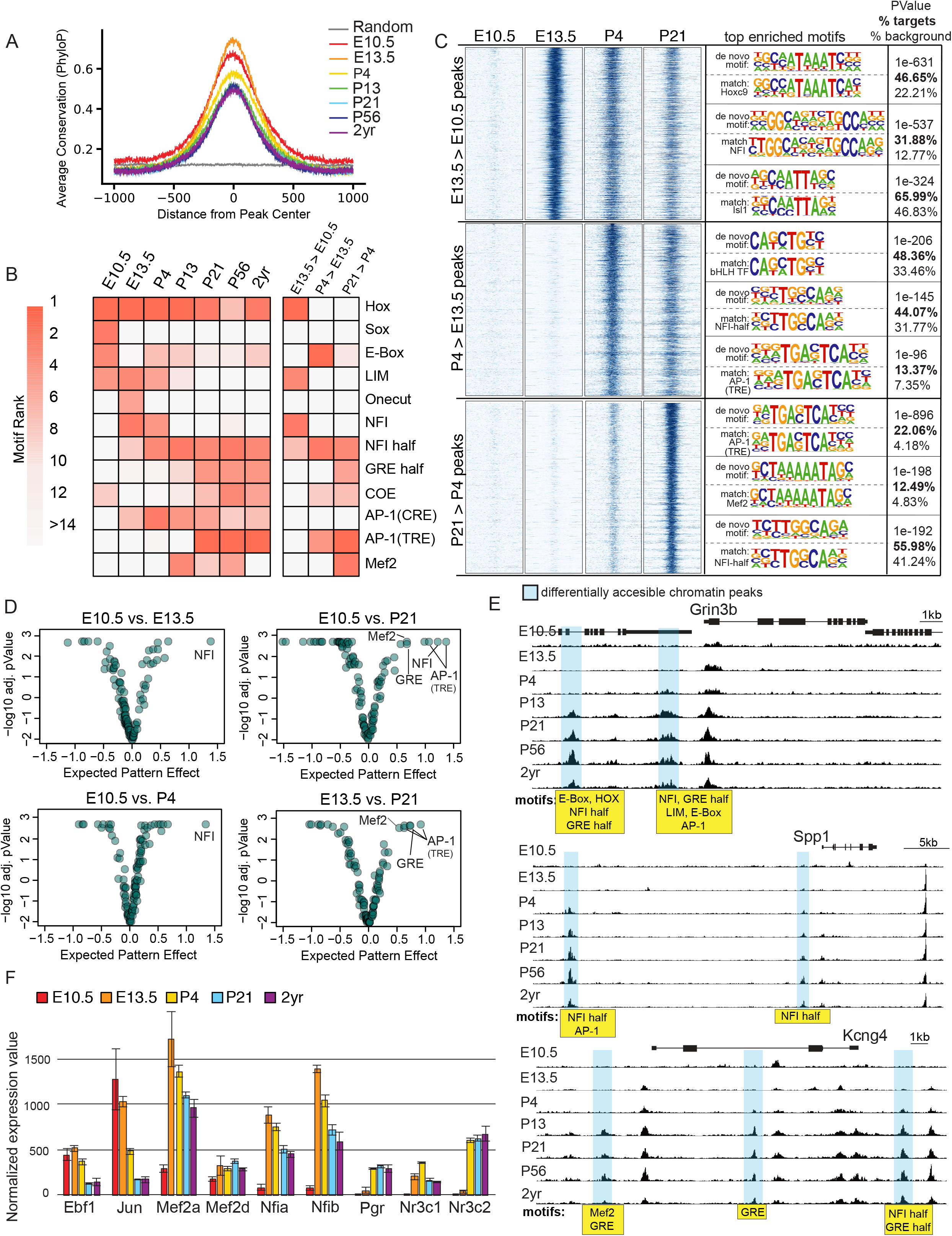
Motif enrichment in age-specific accessible regions identifies putative regulators of motor neuron maturation. A) The conservation of motor neuron accessible regions at each age compared to random genomic sequences. The motor neuron accessible sites were generated by filtering out accessible regions in ESCs and CTCF bound sites, and then taking the top 10k most significant ATAC-seq peaks. Random sequences are ~50k randomly sampled 2kb regions from the genome. B) Heatmap of the most highly enriched motifs. The left panel shows the most highly enriched motifs in the most significant 10k regions at each age (same as A). The right panel shows the most highly enriched motifs in the most significant 10k differentially accessible peaks (same as C). For example, peaks that are not accessible at E10.5, but become accessible at E13.5 (E13.5 > E10.5), etc. C) Motifs enriched in differentially accessible regions. Left: Heatmaps showing ATAC-seq reads (counts per million) at 10k most significant genomic regions that are differentially accessible with age. Each panel spans +/− 1kb from the center of peak. Middle: the top three de novo motifs enriched within the differentially accessible regions and the known transcription factor motif that best matches the de novo motif. Right: The pValue and prevalence of enriched motif, as determined by HOMER. D) Identification of motifs enriched during maturation using DeepAccess. Each dot in the plots represent one of 108 consensus transcription factor binding motifs. The Expected Pattern Effect (x-axis) is a score that represents how differentially predictive each motif is of accessibility between two timepoints. For example, NFI motifs are highly predictive of accessibility at P4 and E13.5 as compared to E10.5 (left two plots). E) Genome browser views of ATAC-seq peaks around three postnatally upregulated genes. Differentially accessible regions are highlighted in blue and the motifs present in each of these regions are listed in a yellow box. F) Normalized expression values of transcription factors that bind to enriched motifs. Ebf1 binds to COE motifs. Jun binds to AP-1(TRE) motifs in a complex with Fos. Mef2a/d bind to Mef2 motifs when activated. Nfia/b bind to NFI motifs. Pgr, Nr3c1, and Nr3c2 bind to hormone receptor motifs (GRE) in the presence of ligand.

De novo motif enrichment analysis (**Fig. 4b, 4c**) revealed several transcription factor families that are already known to regulate gene expression in motor neurons. First, the LIM motif is highly enriched at E10.5 and the Onecut motif is highly enriched at E13.5. This is in accordance with previous findings that Isl1 regulates gene expression in motor neurons by first binding to LIM motifs with Lhx3, and then binding to Onecut motifs with Onecut1 (Rhee et al., 2016). Secondly, the other top motifs enriched at E10.5 are Sox and E-box. Sox and bHLH transcription factors were previously shown to be expressed in motor neuron progenitors and in nascent motor neurons respectively (Delile et al., 2019; Graham et al., 2003; Mazzoni et al., 2013). Third, the HOX motif is highly enriched at all ages, in agreement with the fact that Hox transcription factors are required for establishing rostro-caudal identity in spinal cord motor neurons (Dasen and Jessell, 2009). The rediscovery of these previously known transcriptional regulators suggests that motif enrichment analyses of accessible regions will be an effective method for discovering age-specific regulators of motor neuron gene expression.

Having validated the approach, we turned to the analysis of motifs enriched in maturing motor neurons. We identified motifs of 5 transcription factor families that were highly enriched in postnatal motor neurons: NFI, Ebf (bind to COE motifs), steroid hormone receptors (bind to GREs), AP-1 (TRE and CRE motifs), and Mef2. NFI motifs become highly enriched at E13.5 and remain enriched at all ages after. Interestingly, NFI motifs are not only enriched in the most significant accessible regions at each age, but are also enriched in chromatin regions that become differentially accessible at successive developmental timepoints (i.e. E13.5 > E10.5, P4 > E13.5, and P21 > P4 accessible regions) (**Fig 4b, 4c**). This suggested that at each subsequent age, these transcription factors bind to a new set of putative regulatory regions. Indeed, when we search for secondary motifs, we find that NFI containing peaks at E13.5 are enriched for HOX and LIM motifs, whereas NFI containing peaks at P21 are enriched for HOX, AP-1, and Mef2 motifs. NFI factors have also been shown to sequentially control maturation events in cerebral granule neurons (Ding et al., 2016; Kilpatrick et al., 2012). Ebf-binding COE motifs are also similarly, but less consistently, enriched at all ages. This transcription factor family has already been shown to play a role in the specification and maintenance of the motor neuron gene program (Kratsios et al., 2011; Velasco et al., 2017). AP-1 and Mef2 motifs are bound by activity dependent transcription factor families. Among the AP-1 motifs, we find the TRE motif which is bound by Fos/Jun transcription factors to be more highly and specifically enriched in postnatal motor neurons; AP-1 was recently shown to also control postnatal gene expression in cortical interneurons (Stroud et al., 2020). The enrichment of steroid hormone receptor motifs in postnatal motor neurons is intriguing as progesterone and mineralocorticoid receptors have been shown to play a role in morphological and functional maturation of purkinje cells and specification of hippocampal neurons (McCann et al., 2021; Tamura et al., 2011; Wessel et al., 2014).

As a secondary approach for motif enrichment, we used DeepAccess, an ensemble of convolutional neural networks which takes as input the DNA sequence from 100nt regions of the genome and is trained to predict accessibility from ATAC-seq data (Hammelman et al., 2020). It then computes the effects of known transcription factor motifs on differential accessibility between two different datasets. In contrast to HOMER, which performs motif enrichment by comparing a positive set of sequences to random genomic sequences, DeepAccess learns the relationship between DNA sequence and accessibility across many datasets, in our case ATAC-seq from mouse ESCs and E10.5-2yr motor neurons. Then a model interpretation technique called Differential Expected Pattern Effect (Hammelman and Gifford, 2021) is used to identify DNA sequence patterns, such as transcription factor motifs, that are differentially predictive of accessibility in one dataset compared to another within the DeepAccess model. When we compared accessibility at E10.5 to E13.5 or E10.5 to P4, we found that among a consensus set of 108 transcription factor motifs tested, the NFI motif is the strongest differential motif distinguishing E13.5 and P4 ages from E10.5. When comparing E10.5 to P21, NFI, AP-1, Mef2, and GRE motifs were highly enriched at P21. Finally, comparing E13.5 to P21, showed the AP-1, Mef2, and GRE motifs most strongly distinguish accessibility at P21 from E13.5, reinforcing the finding that NFI motifs are important for accessibility at both these ages (**Fig. 4d**). Both types of motif enrichment analyses therefore consistently recovered the same significant motifs. A closer look at the genomic regions around three postnatally upregulated genes, Grin3b, Kcng4, and Spp1 (**Supplementary Fig. 1**) shows the presence of several distal regions that are specifically accessible at postnatal ages. Each of these peaks contains at least one transcription factor binding motif that is enriched during maturation (**Fig. 4e**).

We next asked if the transcription factors that bind to the discovered motifs are expressed in motor neurons at the correct timepoints (**Fig. 4f**). We find that this is indeed the case: Ebf1 is expressed at all ages; as previously reported, NFI transcription factors Nfia and Nfib are activated in motor neurons between E10.5 and E13.5 and continuously maintained; Mef2a/d are also continuously maintained after upregulation at E13.5; steroid hormone receptor transcription factors Pgr and Nr3c2 are specifically activated only in postnatal motor neurons and then continuously maintained. The expression of AP-1 binding transcription factor Fos, however, is not very robust at any age, even though Jun expression is detectible at all ages. This is consistent with the fact that Fos is an immediate early gene, expressed only transiently in response to neuronal activation (Yap and Greenberg, 2018). Overall, motif enrichment analysis of accessible chromatin identified several novel candidate transcriptional regulators of motor neuron maturation.

### Genes upregulated during maturation and temporally-dynamic chromatin regions are largely motor neuron specific

Recent gene expression analyses performed in cortical neurons, somatosensory neurons, and hypothalamic neurons demonstrate that many, if not all, post mitotic neurons undergo a protracted period of transcriptional maturation (Romanov et al., 2020; Sharma et al., 2020; Stroud et al., 2020). To determine which aspects of the motor neuron maturation program are cell-type-specific vs. shared with other neurons, we compared gene expression in motor neurons to published data from postnatal cortical neurons (Stroud et al., 2020). The published cortical data is comparable to motor neurons as it was generated by performing bulk RNA-seq of nuclei from VIP and SST inhibitory interneurons and RORB expressing excitatory neurons from postnatal ages P7 to P56. We processed these data using the same methods as motor neurons (Methods) to minimize methodological bias. To identify any broadly-shared aspects of the neuronal maturation program, we identified genes that are up- or down-regulated in motor neurons between P4 and P56 (992 up, 1973 down genes; p-value < 0.001, > 2x fold change), and asked what percent of these genes are also regulated in the same direction in VIP, SST, and RORB neurons (**Supplementary Fig. 2d – 2h**). We found that 12% of genes upregulated in motor neurons are also upregulated in VIP, SST, and RORB neurons, and 44% of the genes downregulated in motor neurons are also downregulated in VIP, SST, and RORB neurons. This shows that whereas the upregulated gene expression program is largely cell type specific, part of the downregulated gene expression program is shared between diverse neuronal types. When we perform pathway enrichment analysis (Jassal et al., 2020) on the shared down-regulated genes, the most enriched pathways are those related to protein metabolism, such as Eukaryotic Translation Initiation (51 genes, p-value 1.11e-16), and pathways related to neural development such as axon guidance (112 genes, p-value 1.11e-16) and signaling by ROBO receptors (57 genes, p-value 3.44 e-15). The shared upregulated genes, on the other hand, were not enriched for any functional pathways using the same analysis method at a threshold p-value less than 0.05.

We performed a complementary analysis of genomic regions that gain or lose accessibility in mature neurons. To identify how many gained accessible regions are shared, we took the top 100k most significant adult peaks in all cell types and subtracted peaks found in P4/P7 animals. This gave us 32,040 adult-specific peaks in SST, 41,388 in VIP, 30,334 peaks in RORB neurons and 40,832 peaks in motor neurons. Remarkably, of these differential peaks, only 159 peaks (0.38% of motor neuron peaks) overlap between all four cell types, showing that adult-specific dynamic peaks are largely unique in individual neuron types. Performing the same analysis on peaks that lose accessibility between P4/P7 and P56, we found 42,524 P7-specific peaks in SST, 52,462 in VIP, 31,895 peaks RORB neurons and 35,385 P4-specific peaks in motor neurons. Only 140 peaks (0.39% of motor neuron peaks) overlap between all four cell types. Even though the down-regulated gene expression program is partially shared between cell types, dynamic regulatory regions are highly cell type specific.

### AP-1, Mef2, and NFI transcription factors are shared regulators of adult identity

We next wanted to understand regulatory mechanisms that control shared and motor neuron specific aspects of adult identity. To do this, we analyzed accessible genomic regions in adult neurons, regardless of whether they are dynamically controlled during maturation. We performed de novo motif enrichment analysis in top shared and neuron-specific accessible chromatin regions in adult motor neurons, VIP and SST interneurons, and RORB excitatory neurons. To identify top accessible regions shared between motor neurons and cortical neurons, we took the top 100k most significant motor neuron accessible regions at P56, filtered out ESC accessible regions and CTCF-bound sites, and then asked how many of the remaining 75,894 peaks were shared with cortical neurons. We found that 19,589 (25%), 18,831(24.8%), and 18,249 (24%) of motor neuron peaks were shared with SST, VIP, and RORB neurons respectively, and 9,337 (12%) peaks have regions that were accessible in all four cell types. ~90% of the shared peaks overlap with regions of H3K27ac in either SST, VIP, or RORB neurons, showing that a majority of shared accessible regions are active enhancers. A similar degree of overlap in adult accessible regions has previously been reported (Mo et al., 2015). It is of interest that the shared and motor neuron-specific peaks are similarly conserved, while SST, VIP, and RORB specific regions exhibit lower degree of conservation (**Fig. 5a**), consistent with the view that motor neurons are among the evolutionarily oldest neuronal cell types (Jung and Dasen, 2015).

**Figure 5:**
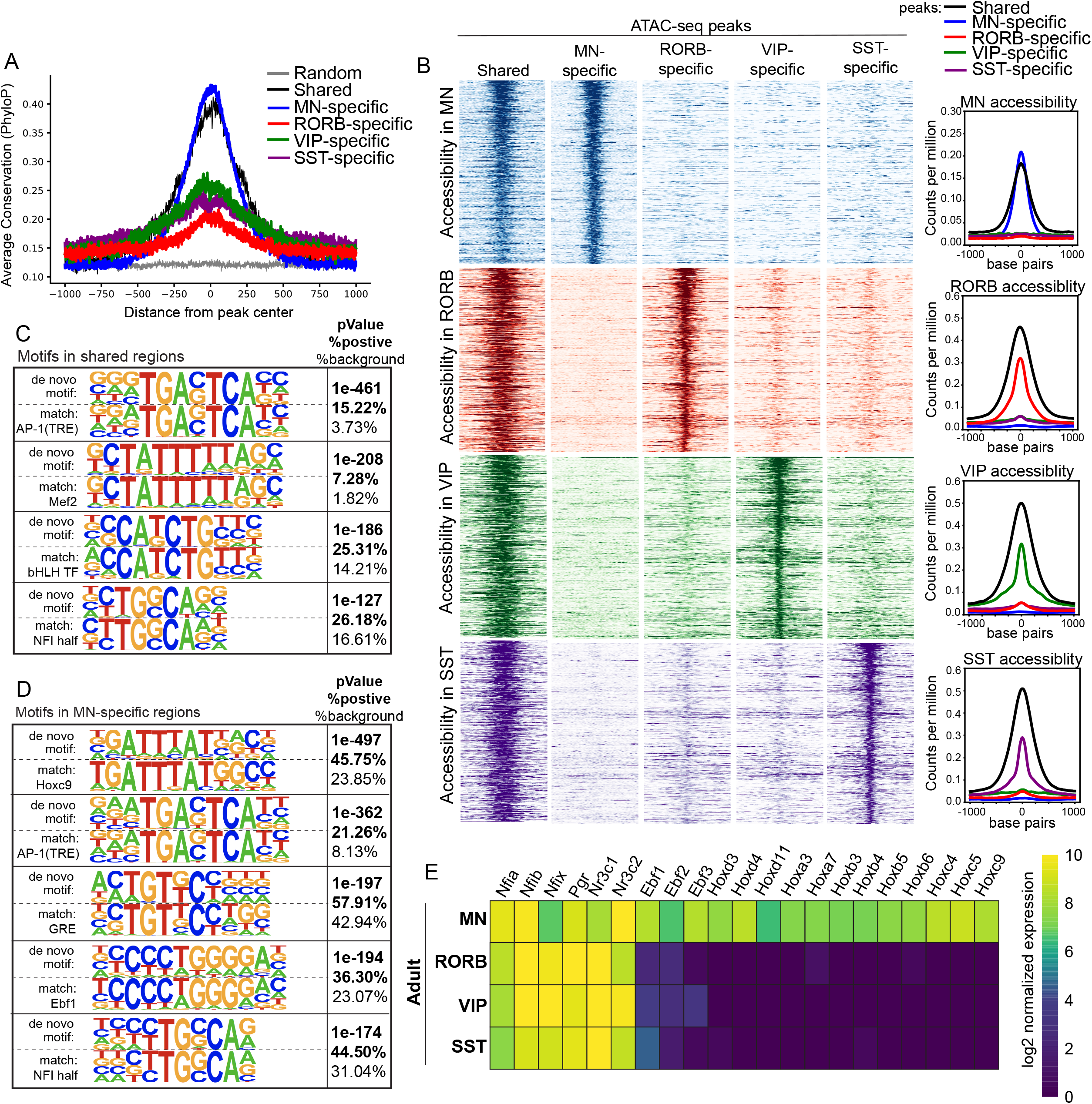
Identification of shared and motor neuron specific regulators of adult gene expression. A) The conservation of shared and neuron-specific adult accessible regions compared to random genomic sequences. Cell type specific regions were determined by taking the most significant 100k ATAC-seq regions in each adult cell type and filtering out the regions that were accessible in any of the other three cell types. Shared regions were those that are present in the 100k most significant accessible regions in all four adult cell types, with ESC-accessible sites and CTCF-binding sites filtered out. Random sequences are ~50k randomly sampled 2kb regions from the genome. B) Heatmaps (left) and summary line plots (right) showing ATAC-seq reads (counts per million at cell type specific and shared accessible regions. Each panel spans +/− 1kb from the center of peak. C) The most highly enriched motifs in shared accessible regions. HOMER de novo motif enrichment analysis was performed on the 10k most significant shared accessible genomic regions. Known transcription factor binding motifs that best match the de novo motifs are also shown along with pValue and prevalence of motif. D) The most highly enriched motifs in motor neuron specific accessible regions. HOMER de novo motif enrichment analysis was performed on the 10k most significant motor neuron specific accessible genomic regions. Known transcription factor binding motifs that best match the de novo motifs are also shown along with pValue and prevalence of motif. E) Heatmap showing the normalized expression values of transcription factors that bind to motor neuron specific accessible regions in four adult cell types.

To identify potential motor neuron specific and shared transcriptional regulators, we performed motif enrichment analysis on the identified sets of accessible regions (**Fig. 5b**). Homer de novo enrichment on shared peaks revealed AP-1, Mef2, E-box and NFI motifs to be the most highly enriched (**Fig. 5c**). By contrast, motor neuron specific peaks are enriched for HOX, Ebf, Hormone Receptor, AP-1, and NFI motifs (**Fig. 5d**). Interestingly, we also note that Hox and Ebf transcription factors are expressed in motor neurons, but not in VIP, SST, or RORB neurons (**Fig. 5e**). This analysis suggests that activity dependent AP-1 and Mef2 transcription factors, and NFI factors are broadly involved in gene regulation in adult neurons. In addition to these shared factors, motor neurons utilize cell-type-specific transcription factors to regulate adult gene expression.

## Discussion

We have characterized gene expression and chromatin accessibility changes that accompany the functional maturation of motor neurons, and performed thorough motif enrichment analysis on regulatory regions to discover motor neuron specific and shared regulators of maturation. This study also sheds light on the temporal dynamics of motor neuron subtype diversification. In combination with other recent studies that map gene expression changes in cortical neuron types, these data serve to illuminate the general principles of molecular maturation in the nervous system.

Even though motor neurons acquire their postmitotic identity during embryonic development, gene expression and chromatin accessibility remain dynamic until the third postnatal week. After this time, the transcriptional identity of motor neurons becomes strikingly stable for the remainder of life. Interestingly, a similar timeline is observed in temporal gene expression studies performed on the whole cortex and in the VIP, SST, and RORB cortical neurons (Jacko et al., 2018; Stroud et al., 2020). These neuron types are both regionally and functionally distinct, suggesting a shared timeline of maturation for the entire nervous system.

We found that a majority of the genes that are specifically expressed in alpha and gamma subtypes of motor neurons are temporally induced during maturation. This finding suggests that motor neurons are refining their functional identities well into postnatal life. In mouse models of ALS, alpha and gamma motor neurons show differential susceptibility to degeneration - while alpha motor neurons are susceptible to degeneration, gamma motor neurons are spared (Lalancette-Hebert et al., 2016). The later activation of alpha vs. gamma identities may contribute to the adult-specific onset of motor neuron degeneration in ALS, as young postnatal motor neurons may not yet have acquired the features that make them vulnerable.

We identified AP-1, Mef2, and NFI factors as shared regulators of maturation. AP-1 and Mef2 factors are activated in response to neuronal activity and other stimuli such as growth factors (Greenberg and Ziff, 1984; Yap and Greenberg, 2018). AP-1 factors have recently been shown to control maturation specific gene expression profiles by binding to maturation specific enhancers in VIP and SST cortical neurons (Stroud et al., 2020). Our analysis shows that binding sites for these factors are enriched in both shared and motor neuron specific accessible regions. This supports the idea that AP-1 factors are part of a core transcriptional program that universally regulates mature gene expression programs in the nervous system by binding to both shared and cell-specific regulatory elements. NFI factors also play broad roles in nervous system development. In cerebellar granule neurons they are required for multiple steps of neuronal development and maturation including proper axon outgrowth, cell migration, dendrite and synapse formation (Ding et al., 2016; Kilpatrick et al., 2012). In the olfactory epithelium, they are involved in the correct specification of late born olfactory receptors (Bashkirova et al., 2020). They are required for the correct specification of late born neuron types in various other regions including the dorsal spinal cord (Sagner et al., 2020). And, they have been shown to accelerate glial competency in human included pluripotent stem cells (Tchieu et al., 2019). In motor neurons, these factors are activated shortly after specification, and remain continuously expressed thereafter (Delile et al., 2019; Deneen et al., 2006; Matuzelski et al., 2020). We found the NFI binding motif to be one of the most enriched motifs at all ages after E13.5. In immature E13.5 neurons, NFI-containing peaks were co-enriched for HOX and LIM motifs while in mature P21 neurons, NFI-containing peaks were co-enriched for HOX, AP-1, and Mef2 motifs. Moreover, our analysis shows the enrichment of NFI binding sites in both shared and cell-specific accessible regions in adult neuron types. Cumulatively, these data suggest that NFI factors bind to both cell type- and stage-specific regulatory regions, likely with different co-factors, to regulate gene expression. It will be interesting to continue to dissect how this transcription factor family regulates such a diverse array of regulatory events in each cell type.

Despite the presence of several shared regulators of maturation, both the upregulated gene expression program, and the dynamic regulatory regions that control maturation seem to be largely cell type specific. Only ~0.4% of postnatally dynamic accessible regions are shared between motor neurons, VIP interneurons, SST interneurons, and RORB excitatory neurons. These data suggest that shared regulators control cell-specific maturation programs by binding largely distinct regulatory regions in each cell type. In motor neurons, cell type specificity may be achieved by the cooperative action of motor neuron specific and shared regulators. Consistent with this hypothesis, we identified COE, HOX, and GRE motifs in addition to NFI and AP-1 motifs in motor neuron specific, adult accessible regions. The COE transcription factors Ebf1/2/3 and several Hox transcription factors are selectively expressed in motor neurons. These transcription factor families play well established roles in the specification of mouse motor neurons (Dasen and Jessell, 2009; Velasco et al., 2017). The *C. elegans* COE transcription factor *unc-3* is known to be required for both specifying and stably maintaining motor neuron identity (Kratsios et al., 2011; Velasco et al., 2017). Our analysis suggests that Ebf and Hox factors may be continuously required to maintain mouse motor neuron identity, and to direct the cell type specific activity of shared maturation regulators. Together, the data and analyses in this study identified a general transcriptional program that is deployed in a highly cell type- and cell stage-specific manner to control neuronal maturation.

The ability to produce motor neurons *in vitro* from mouse and human stem cells holds great promise for dissecting neuronal function and dysfunction in disease. However, the immature state of stem cell derived neurons makes them an unreliable model for late-onset diseases (Arbab et al., 2014; Bucchia et al., 2018). In this work, we have mapped out the detailed trajectory of motor neuron maturation and identified potential shared and cell-type-specific regulators. These insights could potentially be employed to transcriptionally program the age of stem cell derived motor neurons with a goal to better understand the pathology of age-specific motor neuron diseases.

## Methods

### Mice

Animals were handled according to protocols approved by the Columbia Institute of Comparative Medicine. Mice were housed in a pathogen-free barrier facility with a 12-hour light/dark cycle. All mouse lines used were in pure C57BL/6J background and have been previously characterized: SUN1-sfGFP-Myc (Mo et al., 2015), ChAT-IRES-Cre::SV40pA::Δneo (Chat-Cre) (Rossi et al., 2011), and Hb9-GFP transgenic mice (Wichterle et al., 2002). For purification of whole cells at E10.5, heterozygous Hb9-GFP males were crossed to wildtype C57BL/6J females, and heterozygous Hb9-GFP embryos were sacrificed for motor neuron purification. 3-5 embryos were used per replicate for these experiments. For motor neuron nuclei purification at all other ages, homozygous SUN1-2xsfGFP-6xMYC^1^ mice were crossed to homozygous ChAT-IRES-Cre::SV40pA::Δneo mice and transheterozygous progeny were sacrificed for experiments. For each replicate of RNA-seq and ATAC-seq experiment at each age, spinal cord tissue was combined from 4-7 animals. Both males and females were included in each experiment. Three replicates of RNA-seq experiments were performed at E10.5, P4, P21, and P56, and two replicates were performed at E13.5, P13, and 2yr. Two replicates of ATAC-seq were performed at all ages.

### Immunostaining of spinal cord tissue

#### E10

brachial, thoracic, lumbar vertebral columns were dissected from embryos and fixed in 4% paraformaldehyde for 2 hours. Fixed vertebral columns were washed in PBS 3x times for 10min, 30min, and 1hr. Spinal cords were carefully dissected out of vertebral columns and cryopreserved in OCT. P21: Deeply anesthetized mice were transcardially perfused with 20 mls of PBS, followed by 20mls of 4% paraformaldehyde. Vertebral columns were dissected out and fixed overnight in 4% paraformaldehyde at 4C, followed by 3 PBS washes lasting 1 hr, 2 hr, and overnight. Brachial, thoracic, lumbar spinal cords were dissected out of the vertebral columns and cryopreserved in OCT.

#### Immunostaining

Cryopreserved spinal cords were sectioned into 15um slices onto slides. Sections were blocked for 30 min in Blocking Buffer (10% donkey serum (EMD Millipore), 0.2% Triton X-100 (Sigma-Aldrich) in 1X PBS, and 0.05% NaN3), followed by primary antibody treatment in Antibody Buffer (2% donkey serum, 0.2% Triton X-100 in 1X PBS and 0.05% NaN3), followed by 3x 5min washes with Wash Buffer(0.1% Triton X-100 in PBS), followed by secondary antibody treatment in Antibody Buffer for 2 hours at room temperature, followed by 1x wash with Wash Buffer + DAPI for 15 min, and 2x washes with just Wash Buffer for 5 min each. Sections were washed one last time with PBS, and sealed with Flouromount G (Thermo Fisher Scientific OB100-01) and coverslips. The following primary antibodies were used: GFP (Chick, 1:3000), Chat (Goat, EMD Millipore AB144P 1:100), hb9 (Guniea pig 1:100 from Jessell Lab), NeuN (Rabbit, Millipore Sigma ABN78, 1:1000), Spp1 (Mouse, R&D Systems AF808, 1:50). Primary antibody incubations were done overnight at 4C except for Chat, which was incubated at room temperature for 48 hours. The following secondary antibodies were used: 1:800, Cy3 and Alexa488 dyes, Jackson Immunoresearch Laboratories.

### Imaging

Images were acquired with 20X, 40X oil, 60X oil objectives using confocal laser scanning microscope (LSM Zeiss Meta 510 or 780). Images were processed offline using Image-J.

### Purification of whole motor neurons from Hb9-GFP mice for RNA-seq and ATAC-seq

Brachial, thoracic, lumbar spinal cords were dissected out of E10.5 embryos in HBSS buffer. Spinal cords were cut into small pieces, pipetted into 1.5ml tubes, and pelleted with a brief spin. Spinal cord tissue was then resuspended in Accumax for dissociation. 1ml of Accumax (Sigma-Aldrich A7089-100ML) with 5ul of 5mg/ml DNAseI (Millipore-Sigma DN25-10MG) was used for 2 spinal cords. Dissociation was performed by incubating tubes at 37C while shaking at 1000rpm for 15 min. Tissue was then homogenized by pipetting up and down ~20 times, and mixed with 3ml of L15 media (with 1:200 5mg/ml DNAseI). The resulting cell suspension was spun down at 300g for 5 min to pellet single cells. Cells were resuspended in Sorting Buffer (Ca/Mg free PBS with 1% FBS), and filtered through a 20um filter. High GFP cells were sorted using a BioRad Se3 sorter into Sorting Buffer. For RNA collection, cells were spun down, resuspended in TRIzol (Life Technologies 5596-018), flash frozen, and stored at −80C till RNA purification. For ATAC-seq, cells were spun down and ATAC-seq was performed exactly as published in (Buenrostro et al., 2015).

### INTACT nuclear isolation for RNA-seq

We used a modified version of the protocol in (Mo et al., 2015) with all the same chemical reagents. The following volumes are for collecting nuclei from 3-4 spinal cords from E13.5/P4 mice or 2 spinal cords from P13 or older mice. Volumes have to be adjusted for higher numbers of spinal cords being processed together. Brachial and lumbar regions of the spinal cords were dissected out and processed together for RNA-seq. Brachial, thoracic, and lumbar regions of the spinal cords were dissected and processed together for ATAC-seq.

#### Purification of nuclei

Spinal cords were homogenized by douncing 10x times (E13.5, P4) or 15-20x times (P13, P21, P56, 2yr) with a 1ml dounce-homogenizer in a total of 5ml Buffer HB (0.25M sucrose, 25mM KCl, 5mM MgCl2, 20mM Tricine-KOH pH7.8, 1mM DTT, 0.15 mM spermine, 0.5 mM spermidine, Roche protease inhibitor tablets) supplemented with 1.5ul/ml RNAse Inhibitor (RNasin Plus RNase Inhibitor, Promega N2615) and 0.3% IGEPAL CA-630. Homogenized spinal cords were filtered through a 100um strainer and combined with 1:1 volume of 50% iodixanol (5 volume of OptiPrep[60% iodixanol] + 1 volume of Diluent [150 mM KCl., 30 mM MgCl2, 120 mM Tricine-KOH pH 7.8]), creating a 10ml suspension that is 25% iodixanol. The 25% iodixanol suspension was split equally into two 15ml falcon tubes, such that each tube had 5ml. The 5ml 25% iodixanol suspension in each 15ml tube was underlaid with 2ml of 40% iodixanol (50% iodixanol solution diluted in Buffer HB). Tubes were spun at 1000g for 12 min at 4C with slow acceleration and deceleration (set to ‘3’). After the spin, a layer of cellular debris collects on top of the 25% suspension. This was cleared off by pipetting. 1.2ml of nuclei were then pipetted from the interface of 25% and 40% layers from each 15ml tube. The nuclei were dounce-homogenized 5x times in a 1ml dounce-homogenizer. This step is necessary for eliminating most of the clumps that form when nuclei stick together during the spin, and increases the final purity of Sun1-GFP+ nuclei.

#### Affinity purification of GFP+ nuclei

1.2ml of nuclei were pipetted into a 1.5ml tube. 70 ul of ProteinG Dynabeads (Life Technologies, 100004D) were washed 2x times with 800ul wash buffer (0.25M sucrose, 25mM KCl, 5mM MgCl2, 20mM Tricine-KOH pH7.8, 0.4% IGEPAL CA-630, 1mM DTT, 0.15 mM spermine, 0.5 mM spermidine) and returned to their original volume (70ul) in wash buffer. 10ul of washed ProteinG beads were added to 1.2ml of nuclei for preclearing, for which tubes were incubated at 4C for 30min with rotation. At this point, the material bound to the beads is non-specific and needs to be removed from solution. To do this, tubes were placed on a magnet for 2 min and 1.2ml of precleared nuclear suspension was then pipetted into a new 1.5ml tube (leaving behind preclearing beads, which can be discarded). For IP reaction, the following was then added to each tube containing 1.2ml of nuclei: 5ul anti-GFP antibody (Thermo Fisher Scientific G10362), 200 ul Wash Buffer, 1ul RNase Inhibitor. The IP reactions were incubated for 30 minutes at 4C with rotation. After incubation, 25ul pre-washed beads were added to each IP reaction, followed by an additional 20 minutes of rotation at 4C. At this point the samples contain beads that are bound to GFP+ nuclei. To increase the number of beads bound to each nucleus the following was repeated 6-7 times: tubes were placed on a magnet for no more than 1 min, then put on ice for ~15s, then beads were resuspended into solution by gentle inversion. The tubes were then put on a magnet for 2 min and the liquid suspension (which contains mainly GFP- nulcei) was pipetted off. The beads from all tubes that came from the same starting sample were combined and resuspended in 700ul of wash buffer and filtered through a 20um filter to eliminate large clumps of nuclei. The beads were then washed 5x times by doing the following: tubes were placed on magnet for 1min, Wash Buffer was pipetted off, 1.4ml of fresh Wash Buffer was added, tubes are placed on ice for ~15s, and beads were resuspended by rigorously pipetting. After the last wash, beads were resuspended in 200-300ul of Wash Buffer. 1.5ul of beads were mixed with 1.5ul of DAPI for visualization with fluorescent microscope and determination of purity. Finally, the tubes were placed on magnet, Wash Buffer was removed, beads were resuspended in 500ul of Trizol, and incubated at room temperature for 10min. The tube was then placed on the magnet for a last time, 500ul of TRIzol was pipetted into a new tube which was flash frozen and stored at −80C till RNA purification. This procedure yielded ~10,000 nuclei per spinal cord (using brachial and lumbar sections). At E13.5 and P4, nuclei were ~85% GFP+ with some contamination from blood vessels. At P13 and older ages, nuclei were >95% GFP+.

### FAC sorting nuclei for ATAC-seq

We noticed that bead bound nuclei purified using the above affinity purification method yielded a very low concentration of DNA after ATAC-seq. We therefore FAC sorted nuclei for ATAC-seq experiments. To do this, nuclei were purified from spinal cords following the above method. Then, instead of performing affinity purification, we did the following:

#### FAC sorting of GFP+ nuclei

1.2ml of nuclei were thoroughly mixed with 3.6 ml of Wash Buffer in 5ml Polypropylene Round-Bottom Tube. Tubes were spun at 500g for 5min to pellet nuclei. The liquid suspension was carefully pipetted off. All nuclei from the same starting sample were combined and resuspended in 1.5-2.5 ml of Wash Buffer with rigorous pipetting to ensure no clumps of nuclei remained. The nuclei were then filtered through a 20um filter, and 1.5ul/ml of RNAse inhibitor was added. GFP+ nuclei were then FACs sorted into Wash Buffer with1.5ul/ml of RNAse inhibitor using a BioRad Se3 sorter. 1.5ul of sorted nuclei were mixed with 1.5ul of DAPI to visualize nuclei and determine purity. This procedure resulted in >95% pure nuclei at all ages.

### RNA purification

Tubes containing samples in TRIzol were thawed to room temperature, 100ul of Chloroform was added to 500ul of TRIzol, mixed by rigorous inversion and spun down at 16,000g for 15 min. ~270ul of the aqueous layer was carefully removed. RNA was purified from the aqueous layer using the Zymo RNA microprep kit (ZymoResearch, cat #R2060) following manufacturer’s instructions.

### RNA-seq library preparation and sequencing

RNA-seq libraries were prepared at the MIT BioMicroCenter using the following kit: Clontech SMARTer Stranded Total RNA-Seq Kit – Pico Input Mammalian with ZAPr ribosomal depletion. Paired end sequencing was performed using the 150nt Nextseq kit.

### RNA-seq processing

Reads were trimmed for adaptors and low-quality positions using Trimgalore (Cutadapt v0.6.2) (Martin, 2011). Reads were aligned to the mouse genome (mm10) and gene-level counts were quantified using RSEM (v1.3.0) (Li and Dewey, 2011) rsem-calculate-expression using default parameters and STAR (v2.5.2b) for alignment. Differential gene expression analysis was performed on RSEM gene-level read counts using EdgeR (McCarthy et al., 2012; Robinson et al., 2010). Read counts were normalized by median of ratios normalization (Maza et al., 2013) using a custom Python (v3.6.9) script.

### ATAC-seq on nuclei

FAC sorted nuclei were spun at 500g for 7 min to pellet nuclei. Wash Buffer was carefully pipetted off, leaving behind ~100ul of Wash Buffer. The nuclear pellet was gently resuspended in the remaining 100ul Wash Buffer and transferred to a 0.2ml PCR tube. Nuclei were pelleted again by spinning at 500g for 5min. Almost all of the Wash Buffer was then pipetted off. Nuclei were then resuspended in the tagmentation solution containing Tn5. ATAC-seq was performed using the OMNI-ATAC protocol (Corces et al., 2017). The cell lysis step was skipped as we were starting with nuclei, but the remainder of the protocol was followed exactly as published.

### ATAC-seq library preparation and sequencing

ATAC libraries were amplified by following the published protocol (Buenrostro et al., 2015). All ATAC-seq libraries required 8-9 total amplification cycles. ATAC-seq libraries were sequenced at the MIT BioMicroCenter. Paired end sequencing was performed using the 75nt Nextseq kit.

### ATAC-seq processing

Reads were trimmed for adaptors and low-quality positions using Trimgalore (Cutadapt v0.6.2) (Martin, 2011). Reads were aligned to the mouse genome (mm10) with bwa mem (v0.7.1.7) (Li, 2013) with default parameters. Duplicates were removed with samtools (v1.7.2) (Li et al., 2009) markdup and properly paired mapped reads were filtered. Accessible regions were called using MACS2 (v2.2.7.1) (Zhang et al., 2008) with the parameters -f BAMPE -g mm -p 0.01 --shift −36 --extsize 73 --nomodel -- keep-dup all --call-summits. Accessible regions that overlapped genome blacklist regions were excluded from downstream analysis. The BEDTools (Quinlan and Hall, 2010) suite was used to compare peaks between different samples.

### Motif Discovery Analysis

HOMER (Heinz et al., 2010) was used to perform de novo motif enrichment using the command findMotifsGenome.pl with the parameters - size given. The de novo output was manually curated for instances when very similar motifs were found enriched twice in the same condition, only one was reported. For differential motif activity, we trained a DeepAccess model on 3,555,674 100nt regions. Each region is labeled as accessible or inaccessible in each of eight cell types: ESCs, E10.5, E13.5, P4, P13, P21, P56, and P2yr. We define a region as accessible in a given cell type if more that 50% of the 100nt region overlaps a MACS2 accessible region from that cell type. 2,555,674 regions were open in at least 1 cell type, and 1,000,000 regions were closed in all cell types (randomly sampled from the genome). Chromosomes 18 and 19 are held out for validation and testing. Methods for computing Differential Expected Pattern Effects between cell types are described in Hammelman and Gifford, 2021. Briefly, we compute a differential expected pattern effect as the ratio between the effect that the presence of a transcription factor motif has on the predicted accessibility of a DNA sequence in one cell type relative to another cell type within a DeepAccess model. We use a customized consensus database of 108 transcription factor motifs representing the major transcription factor families which we derived from the HOCOMOCOv11 database.

### Conservation Analysis

Positional phastCons and PhyloP mouse conservation scores (Pollard et al., 2010) were downloaded as bigwigs from UCSC. Average per-base conservation scores over bed accessible regions were calculated using a custom Python (v3.6.9) script. For comparison, we compare conservation within accessible regions to the conservation of 49,896 2kb regions randomly sampled from the genome.

### PCA

Using a custom python script, normalized gene counts are log-transformed and filtered to keep only genes with more than 10 normalized reads in at least 1 sample. The gene by sample matrix is mean-centered and scaled to unit variance and used as input to perform PCA. Scaling and PCA was performed in Python (v3.6.9) with the sklearn package.

### Published data used in this study

The cortical RNA-seq and ATAC-seq data from Stroud et al., 2020 was downloaded from Gene Expression Omnibus (GEO): GSE150538.

The ESC ATAC-seq data from Dieuleveult et al., 2016 was downloaded from Gene Expression Omnibus (GEO): GSE64825

Raw data from cortical neurons and ESCs was downloaded and processed as described above.

The snRNA-seq data from Blum et al., 2021 is on GEO: GSE161621. List of alpha and gamma specific genes was taken form Blum et al., 2021 Figure 3d. Short list of pool identity markers was taken from Figure 4c. Long list of 322 alpha pool identity markers was taken from Supplementary Table 2f.

## Author Contributions

T.P. and H.W. conceptualized and designed the project. T.P. performed all experiments. All genomic data was processed by J.H., analyzed by T.P. and J.H. with help from M.C. and feedback from D.G and H.W. T.P. and H.W. wrote the manuscript with input from all authors.

## Data and Code Availability

All data generated in this study will be made available on GEO upon publication. Code used for plotting will be made available on GitHub.

## Acknowledgements

We are very grateful to Aaron Gitler and Jacob Blum for sharing snRNA-seq data from adult spinal cords before publication of their manuscript. We also thank all members of the Wichterle, Gifford, Zhang, and Au labs for feedback on the project. We thank our funding sources: T.P. was funded by NINDS Postdoctoral NRSA Fellowship (F32NS105372). J.H. was funded by NSF Graduate Research Fellowship (1122374). H.W. is funded by R01NS109217-01 and R01NS116141.

## Ethics Declarations

The authors declare no competing interests.

## Supplementary Figure Legends

**Supplementary Figure 1:**
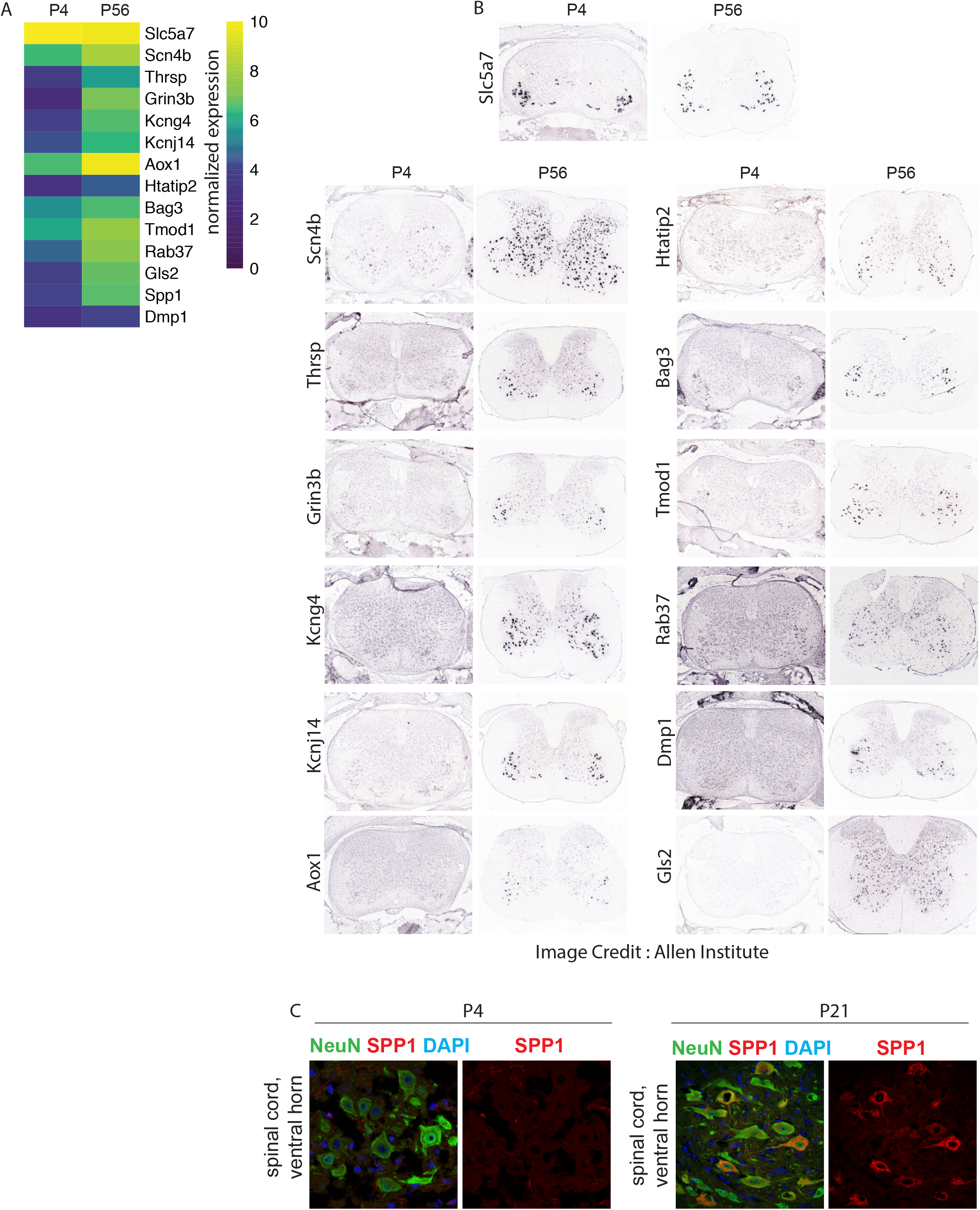
Validation of genes upregulated during motor neuron maturation. A) Heat map showing expression of a continuously expressed gene, Slc5a7, and 13 genes that are upregulated over time. B) Allen Spinal Cord Atlas in situ images of genes from (A) at P4 and P56. C) Immunostaining of OPN, protein encoded by Spp1 gene, in P4 motor neurons and P21 motor neurons. Image shows the ventral horn of the spinal cord.

**Supplementary Figure 2:**
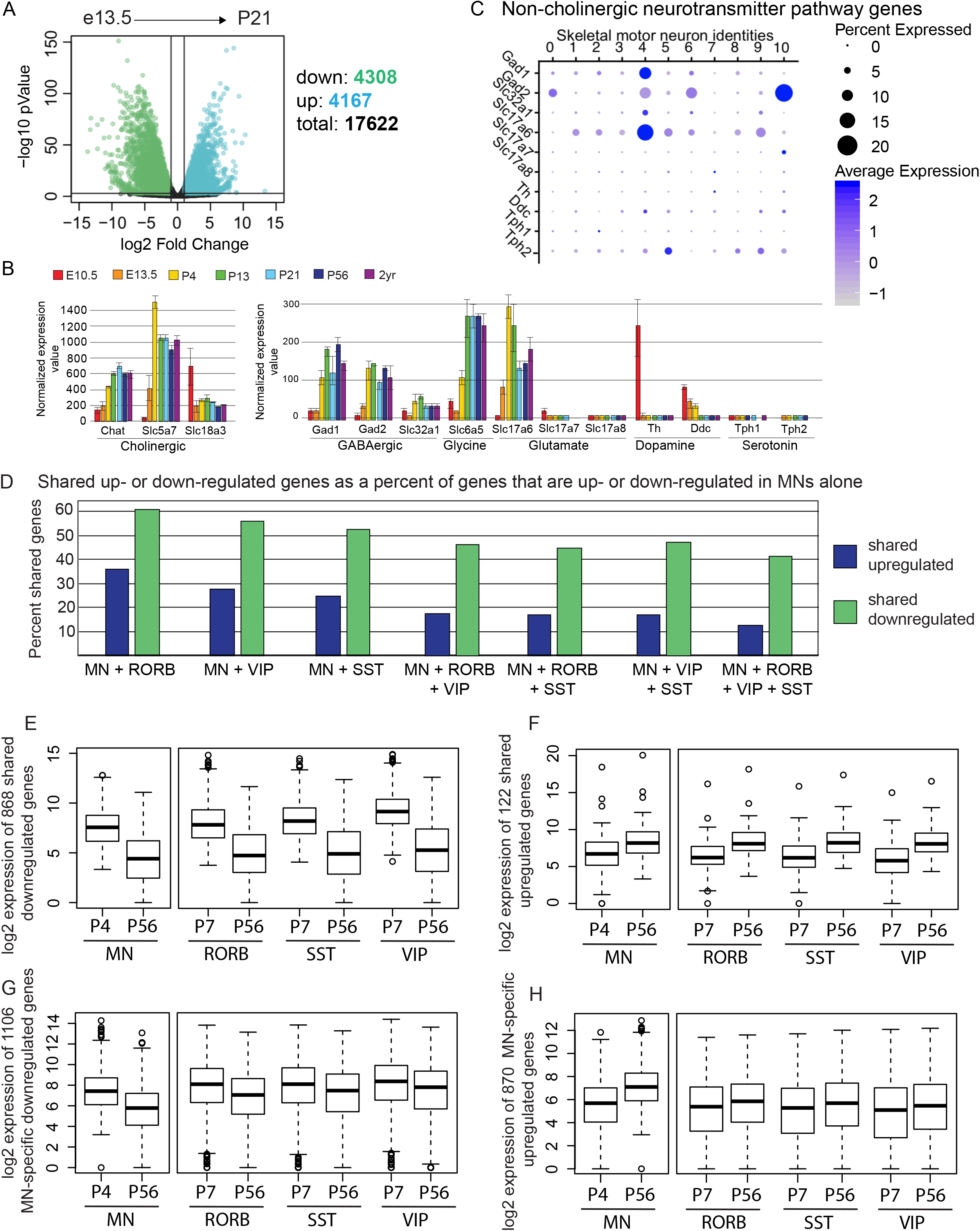
Shared and motor neuron specific changes in gene expression. A) Differential gene expression between E13.5 and P21. Each dot represents one gene, the x-axis shows the log_2_ fold change, the y-axis shows the −log_10_ pValue. Colored dots are genes that are upregulated (blue) or downregulated (green) at least 2-fold with a pValue < 0.001 (as determined by EdgeR). The number of up- and down-regulated genes are listed. B) Normalized expression values of neurotransmitter pathway genes over time. The scale of the y-axis is different in the left and right plots. C) Dot plot showing expression of neurotransmitter pathway genes in snRNA-seq data performed by (Blum et al., 2021). D) Percent of motor neuron up- and down-regulated genes that are shared between cell types. The cell types being compared are listed on the x-axis. For motor neurons, gene expression between P56 and P4 timepoints are compared. For VIP, SST, and RORB neurons, gene expression between P7 and P56 timepoints are compared. E) Log_2_ expression values of shared down-regulated genes. F) Log_2_ expression values of shared up-regulated genes G) Log_2_ expression values of motor neuron specific down-regulated genes. H) Log_2_ expression values of motor neuron specific up-regulated genes

## References

Altman, J., and Sudarshan, K. (1975). Postnatal development of locomotion in the laboratory rat. Anim Behav 23, 896–920.

Arbab, M., Baars, S., and Geijsen, N. (2014). Modeling motor neuron disease: the matter of time. Trends Neurosci 37, 642–652.

Arber, S., Ladle, D.R., Lin, J.H., Frank, E., and Jessell, T.M. (2000). ETS gene Er81 controls the formation of functional connections between group Ia sensory afferents and motor neurons. Cell 101, 485–498.

Bashkirova, E., Monahan, K., Campbell, C.E., Osinski, J.M., Tan, L., Schieren, I., Barnea, G., Xie, X.S., Gronostajski, R.M., and Lomvardas, S. (2020). Homeotic Regulation of Olfactory Receptor Choice via NFI-dependent Heterochromatic Silencing and Genomic Compartmentalization. bioRxiv, 2020.2008.2030.274035.

Bikoff, J.B., Gabitto, M.I., Rivard, A.F., Drobac, E., Machado, T.A., Miri, A., Brenner-Morton, S., Famojure, E., Diaz, C., Alvarez, F.J., et al. (2016). Spinal Inhibitory Interneuron Diversity Delineates Variant Motor Microcircuits. Cell 165, 207–219.

Bishop, B. (1982a). Neural plasticity: Part 1. Plasticity in the developing nervous system: prenatal maturation. Phys Ther 62, 1122–1131.

Bishop, B. (1982b). Neural plasticity: Part 2. Postnatal maturation and function-induced plasticity. Phys Ther 62, 1132–1143.

Blum, J.A., Klemm, S., Shadrach, J.L., Guttenplan, K.A., Nakayama, L., Kathiria, A., Hoang, P.T., Gautier, O., Kaltschmidt, J.A., Greenleaf, W.J., et al. (2021). Single-cell transcriptomic analysis of the adult mouse spinal cord reveals molecular diversity of autonomic and skeletal motor neurons. Nat Neurosci.

Bucchia, M., Merwin, S.J., Re, D.B., and Kariya, S. (2018). Limitations and Challenges in Modeling Diseases Involving Spinal Motor Neuron Degeneration in Vitro. Front Cell Neurosci 12, 61.

Buenrostro, J.D., Wu, B., Chang, H.Y., and Greenleaf, W.J. (2015). ATAC-seq: A Method for Assaying Chromatin Accessibility Genome-Wide. Curr Protoc Mol Biol 109, 21–29 21–21 29 29.

Catela, C., Correa, E., Wen, K., Aburas, J., Croci, L., Consalez, G.G., and Kratsios, P. (2019). An ancient role for collier/Olf/Ebf (COE)-type transcription factors in axial motor neuron development. Neural Dev 14, 2.

Corces, M.R., Trevino, A.E., Hamilton, E.G., Greenside, P.G., Sinnott-Armstrong, N.A., Vesuna, S., Satpathy, A.T., Rubin, A.J., Montine, K.S., Wu, B., et al. (2017). An improved ATAC-seq protocol reduces background and enables interrogation of frozen tissues. Nat Methods 14, 959–962.

Dasen, J.S., De Camilli, A., Wang, B., Tucker, P.W., and Jessell, T.M. (2008). Hox repertoires for motor neuron diversity and connectivity gated by a single accessory factor, FoxP1. Cell 134, 304–316.

Dasen, J.S., and Jessell, T.M. (2009). Hox networks and the origins of motor neuron diversity. Curr Top Dev Biol 88, 169–200.

de Dieuleveult, M., Yen, K., Hmitou, I., Depaux, A., Boussouar, F., Bou Dargham, D., Jounier, S., Humbertclaude, H., Ribierre, F., Baulard, C., et al. (2016). Genome-wide nucleosome specificity and function of chromatin remodellers in ES cells. Nature 530, 113–116.

De Marco Garcia, N.V., and Jessell, T.M. (2008). Early motor neuron pool identity and muscle nerve trajectory defined by postmitotic restrictions in Nkx6.1 activity. Neuron 57, 217–231.

Delile, J., Rayon, T., Melchionda, M., Edwards, A., Briscoe, J., and Sagner, A. (2019). Single cell transcriptomics reveals spatial and temporal dynamics of gene expression in the developing mouse spinal cord. Development 146.

Deneen, B., Ho, R., Lukaszewicz, A., Hochstim, C.J., Gronostajski, R.M., and Anderson, D.J. (2006). The transcription factor NFIA controls the onset of gliogenesis in the developing spinal cord. Neuron 52, 953–968.

Ding, B., Cave, J.W., Dobner, P.R., Mullikin-Kilpatrick, D., Bartzokis, M., Zhu, H., Chow, C.W., Gronostajski, R.M., and Kilpatrick, D.L. (2016). Reciprocal autoregulation by NFI occupancy and ETV1 promotes the developmental expression of dendrite-synapse genes in cerebellar granule neurons. Mol Biol Cell 27, 1488–1499.

Donatelle, J.M. (1977). Growth of the corticospinal tract and the development of placing reactions in the postnatal rat. J Comp Neurol 175, 207–231.

Ericson, J., Thor, S., Edlund, T., Jessell, T.M., and Yamada, T. (1992). Early stages of motor neuron differentiation revealed by expression of homeobox gene Islet-1. Science 256, 1555–1560.

Francius, C., and Clotman, F. (2010). Dynamic expression of the Onecut transcription factors HNF-6, OC-2 and OC-3 during spinal motor neuron development. Neuroscience 165, 116–129.

Graham, V., Khudyakov, J., Ellis, P., and Pevny, L. (2003). SOX2 functions to maintain neural progenitor identity. Neuron 39, 749–765.

Greenberg, M.E., and Ziff, E.B. (1984). Stimulation of 3T3 cells induces transcription of the c-fos proto-oncogene. Nature 311, 433–438.

Guo, Y., Xu, Q., Canzio, D., Shou, J., Li, J., Gorkin, D.U., Jung, I., Wu, H., Zhai, Y., Tang, Y., et al. (2015). CRISPR Inversion of CTCF Sites Alters Genome Topology and Enhancer/Promoter Function. Cell 162, 900–910.

Hammelman, J., and Gifford, D.K. (2021). Discovering differential genome sequence activity with interpretable and efficient deep learning. bioRxiv, 2021.2002.2026.433073.

Hammelman, J., Krismer, K., Banerjee, B., Gifford, D.K., and Sherwood, R.I. (2020). Identification of determinants of differential chromatin accessibility through a massively parallel genome-integrated reporter assay. Genome Res 30, 1468–1480.

Heinz, S., Benner, C., Spann, N., Bertolino, E., Lin, Y.C., Laslo, P., Cheng, J.X., Murre, C., Singh, H., and Glass, C.K. (2010). Simple combinations of lineage-determining transcription factors prime cis-regulatory elements required for macrophage and B cell identities. Mol Cell 38, 576–589.

Hoang, P.T., Chalif, J.I., Bikoff, J.B., Jessell, T.M., Mentis, G.Z., and Wichterle, H. (2018). Subtype Diversification and Synaptic Specificity of Stem Cell-Derived Spinal Interneurons. Neuron 100, 135–149 e137.

Jacko, M., Weyn-Vanhentenryck, S.M., Smerdon, J.W., Yan, R., Feng, H., Williams, D.J., Pai, J., Xu, K., Wichterle, H., and Zhang, C. (2018). Rbfox Splicing Factors Promote Neuronal Maturation and Axon Initial Segment Assembly. Neuron 97, 853–868 e856.

Jassal, B., Matthews, L., Viteri, G., Gong, C., Lorente, P., Fabregat, A., Sidiropoulos, K., Cook, J., Gillespie, M., Haw, R., et al. (2020). The reactome pathway knowledgebase. Nucleic Acids Res 48, D498–D503.

Jessell, T.M. (2000). Neuronal specification in the spinal cord: inductive signals and transcriptional codes. Nat Rev Genet 1, 20–29.

Jung, H., and Dasen, J.S. (2015). Evolution of patterning systems and circuit elements for locomotion. Dev Cell 32, 408–422.

Kanning, K.C., Kaplan, A., and Henderson, C.E. (2010). Motor neuron diversity in development and disease. Annu Rev Neurosci 33, 409–440.

Kaplan, A., Spiller, K.J., Towne, C., Kanning, K.C., Choe, G.T., Geber, A., Akay, T., Aebischer, P., and Henderson, C.E. (2014). Neuronal matrix metalloproteinase-9 is a determinant of selective neurodegeneration. Neuron 81, 333–348.

Kilpatrick, D.L., Wang, W., Gronostajski, R., and Litwack, E.D. (2012). Nuclear factor I and cerebellar granule neuron development: an intrinsic-extrinsic interplay. Cerebellum 11, 41–49.

Klemm, S.L., Shipony, Z., and Greenleaf, W.J. (2019). Chromatin accessibility and the regulatory epigenome. Nat Rev Genet 20, 207–220.

Kratsios, P., Stolfi, A., Levine, M., and Hobert, O. (2011). Coordinated regulation of cholinergic motor neuron traits through a conserved terminal selector gene. Nat Neurosci 15, 205–214.

Lake, B.B., Chen, S., Sos, B.C., Fan, J., Kaeser, G.E., Yung, Y.C., Duong, T.E., Gao, D., Chun, J., Kharchenko, P.V., et al. (2018). Integrative single-cell analysis of transcriptional and epigenetic states in the human adult brain. Nat Biotechnol 36, 70–80.

Lalancette-Hebert, M., Sharma, A., Lyashchenko, A.K., and Shneider, N.A. (2016). Gamma motor neurons survive and exacerbate alpha motor neuron degeneration in ALS. Proc Natl Acad Sci U S A 113, E8316–E8325.

Laub, F., Aldabe, R., Ramirez, F., and Friedman, S. (2001). Embryonic expression of Kruppel-like factor 6 in neural and non-neural tissues. Mech Dev 106, 167–170.

Lee, S., Shen, R., Cho, H.H., Kwon, R.J., Seo, S.Y., Lee, J.W., and Lee, S.K. (2013). STAT3 promotes motor neuron differentiation by collaborating with motor neuron-specific LIM complex. Proc Natl Acad Sci U S A 110, 11445–11450.

Lee, S.K., and Pfaff, S.L. (2003). Synchronization of neurogenesis and motor neuron specification by direct coupling of bHLH and homeodomain transcription factors. Neuron 38, 731–745.

Li, H. (2013). Aligning sequence reads, clone sequences and assembly contigs with BWA-MEM. arXiv preprint arXiv:1303.3997

Li, B., and Dewey, C.N. (2011). RSEM: accurate transcript quantification from RNA-Seq data with or without a reference genome. BMC Bioinformatics 12, 323.

Li, H., Handsaker, B., Wysoker, A., Fennell, T., Ruan, J., Homer, N., Marth, G., Abecasis, G., Durbin, R., and Genome Project Data Processing, S. (2009). The Sequence Alignment/Map format and SAMtools. Bioinformatics 25, 2078–2079.

Martin, M. (2011). Cutadapt removes adapter sequences from high-throughput sequencing reads. EMBnet. journal, 17(1), 10–12. doi:https://doi.org/10.14806/ej.17.1.200

Matuzelski, E., Harvey, T.J., Harkins, D., Nguyen, T., Ruitenberg, M.J., and Piper, M. (2020). Expression of NFIA and NFIB within the murine spinal cord. Gene Expr Patterns 35, 119098.

Maza, E., Frasse, P., Senin, P., Bouzayen, M., and Zouine, M. (2013). Comparison of normalization methods for differential gene expression analysis in RNA-Seq experiments: A matter of relative size of studied transcriptomes. Commun Integr Biol 6, e25849.

Mazzoni, E.O., Mahony, S., Closser, M., Morrison, C.A., Nedelec, S., Williams, D.J., An, D., Gifford, D.K., and Wichterle, H. (2013). Synergistic binding of transcription factors to cell-specific enhancers programs motor neuron identity. Nat Neurosci 16, 1219–1227.

McCann, K.E., Lustberg, D.J., Shaughnessy, E.K., Carstens, K.E., Farris, S., Alexander, G.M., Radzicki, D., Zhao, M., and Dudek, S.M. (2021). Novel role for mineralocorticoid receptors in control of a neuronal phenotype. Mol Psychiatry 26, 350–364.

McCarthy, D.J., Chen, Y., and Smyth, G.K. (2012). Differential expression analysis of multifactor RNA-Seq experiments with respect to biological variation. Nucleic Acids Res 40, 4288–4297.

Mendelsohn, A.I., Dasen, J.S., and Jessell, T.M. (2017). Divergent Hox Coding and Evasion of Retinoid Signaling Specifies Motor Neurons Innervating Digit Muscles. Neuron 93, 792–805 e794.

Mentis, G.Z., Alvarez, F.J., Bonnot, A., Richards, D.S., Gonzalez-Forero, D., Zerda, R., and O’Donovan, M.J. (2005). Noncholinergic excitatory actions of motoneurons in the neonatal mammalian spinal cord. Proc Natl Acad Sci U S A 102, 7344–7349.

Mo, A., Mukamel, E.A., Davis, F.P., Luo, C., Henry, G.L., Picard, S., Urich, M.A., Nery, J.R., Sejnowski, T.J., Lister, R., et al. (2015). Epigenomic Signatures of Neuronal Diversity in the Mammalian Brain. Neuron 86, 1369–1384.

Morisaki, Y., Niikura, M., Watanabe, M., Onishi, K., Tanabe, S., Moriwaki, Y., Okuda, T., Ohara, S., Murayama, S., Takao, M., et al. (2016). Selective Expression of Osteopontin in ALS-resistant Motor Neurons is a Critical Determinant of Late Phase Neurodegeneration Mediated by Matrix Metalloproteinase-9. Sci Rep 6, 27354.

Nishimaru, H., Restrepo, C.E., Ryge, J., Yanagawa, Y., and Kiehn, O. (2005). Mammalian motor neurons corelease glutamate and acetylcholine at central synapses. Proc Natl Acad Sci U S A 102, 5245–5249.

Nord, A.S., Blow, M.J., Attanasio, C., Akiyama, J.A., Holt, A., Hosseini, R., Phouanenavong, S., Plajzer-Frick, I., Shoukry, M., Afzal, V., et al. (2013). Rapid and pervasive changes in genome-wide enhancer usage during mammalian development. Cell 155, 1521–1531.

Ong, C.T., and Corces, V.G. (2014). CTCF: an architectural protein bridging genome topology and function. Nat Rev Genet 15, 234–246.

Pfaff, S.L., Mendelsohn, M., Stewart, C.L., Edlund, T., and Jessell, T.M. (1996). Requirement for LIM homeobox gene Isl1 in motor neuron generation reveals a motor neuron-dependent step in interneuron differentiation. Cell 84, 309–320.

Pollard, K.S., Hubisz, M.J., Rosenbloom, K.R., and Siepel, A. (2010). Detection of nonneutral substitution rates on mammalian phylogenies. Genome Res 20, 110–121.

Price, S.R., De Marco Garcia, N.V., Ranscht, B., and Jessell, T.M. (2002). Regulation of motor neuron pool sorting by differential expression of type II cadherins. Cell 109, 205–216.

Quinlan, A.R., and Hall, I.M. (2010). BEDTools: a flexible suite of utilities for comparing genomic features. Bioinformatics 26, 841–842.

Rhee, H.S., Closser, M., Guo, Y., Bashkirova, E.V., Tan, G.C., Gifford, D.K., and Wichterle, H. (2016). Expression of Terminal Effector Genes in Mammalian Neurons Is Maintained by a Dynamic Relay of Transient Enhancers. Neuron 92, 1252–1265.

Richards, D.S., Griffith, R.W., Romer, S.H., and Alvarez, F.J. (2014). Motor axon synapses on renshaw cells contain higher levels of aspartate than glutamate. PLoS One 9, e97240.

Robinson, M.D., McCarthy, D.J., and Smyth, G.K. (2010). edgeR: a Bioconductor package for differential expression analysis of digital gene expression data. Bioinformatics 26, 139–140.

Romanov, R.A., Tretiakov, E.O., Kastriti, M.E., Zupancic, M., Haring, M., Korchynska, S., Popadin, K., Benevento, M., Rebernik, P., Lallemend, F., et al. (2020). Molecular design of hypothalamus development. Nature.

Rossi, J., Balthasar, N., Olson, D., Scott, M., Berglund, E., Lee, C.E., Choi, M.J., Lauzon, D., Lowell, B.B., and Elmquist, J.K. (2011). Melanocortin-4 receptors expressed by cholinergic neurons regulate energy balance and glucose homeostasis. Cell Metab 13, 195–204.

Sagner, A., Zhang, I., Watson, T., Lazaro, J., Melchionda, M., and Briscoe, J. (2020). Temporal patterning of the central nervous system by a shared transcription factor code. bioRxiv, 2020.2011.2010.376491.

Sharma, N., Flaherty, K., Lezgiyeva, K., Wagner, D.E., Klein, A.M., and Ginty, D.D. (2020). The emergence of transcriptional identity in somatosensory neurons. Nature 577, 392–398.

Shneider, N.A., Brown, M.N., Smith, C.A., Pickel, J., and Alvarez, F.J. (2009). Gamma motor neurons express distinct genetic markers at birth and require muscle spindle-derived GDNF for postnatal survival. Neural Dev 4, 42.

Smith, C.C., Paton, J.F.R., Chakrabarty, S., and Ichiyama, R.M. (2017). Descending Systems Direct Development of Key Spinal Motor Circuits. J Neurosci 37, 6372–6387.

Stroud, H., Yang, M.G., Tsitohay, Y.N., Davis, C.P., Sherman, M.A., Hrvatin, S., Ling, E., and Greenberg, M.E. (2020). An Activity-Mediated Transition in Transcription in Early Postnatal Neurons. Neuron 107, 874–890 e878.

Sunkin, S.M., Ng, L., Lau, C., Dolbeare, T., Gilbert, T.L., Thompson, C.L., Hawrylycz, M., and Dang, C. (2013). Allen Brain Atlas: an integrated spatio-temporal portal for exploring the central nervous system. Nucleic Acids Res 41, D996–D1008.

Tamura, M., Sajo, M., Kakita, A., Matsuki, N., and Koyama, R. (2011). Prenatal stress inhibits neuronal maturation through downregulation of mineralocorticoid receptors. J Neurosci 31, 11505–11514.

Tchieu, J., Calder, E.L., Guttikonda, S.R., Gutzwiller, E.M., Aromolaran, K.A., Steinbeck, J.A., Goldstein, P.A., and Studer, L. (2019). NFIA is a gliogenic switch enabling rapid derivation of functional human astrocytes from pluripotent stem cells. Nat Biotechnol 37, 267–275.

Velasco, S., Ibrahim, M.M., Kakumanu, A., Garipler, G., Aydin, B., Al-Sayegh, M.A., Hirsekorn, A., Abdul-Rahman, F., Satija, R., Ohler, U., et al. (2017). A Multi-step Transcriptional and Chromatin State Cascade Underlies Motor Neuron Programming from Embryonic Stem Cells. Cell Stem Cell 20, 205–217 e208.

Wessel, L., Balakrishnan-Renuka, A., Henkel, C., Meyer, H.E., Meller, K., Brand-Saberi, B., and Theiss, C. (2014). Long-term incubation with mifepristone (MLTI) increases the spine density in developing Purkinje cells: new insights into progesterone receptor mechanisms. Cell Mol Life Sci 71, 1723–1740.

Wichterle, H., Lieberam, I., Porter, J.A., and Jessell, T.M. (2002). Directed differentiation of embryonic stem cells into motor neurons. Cell 110, 385–397.

Yap, E.L., and Greenberg, M.E. (2018). Activity-Regulated Transcription: Bridging the Gap between Neural Activity and Behavior. Neuron 100, 330–348.

Yue, F., Cheng, Y., Breschi, A., Vierstra, J., Wu, W., Ryba, T., Sandstrom, R., Ma, Z., Davis, C., Pope, B.D., et al. (2014). A comparative encyclopedia of DNA elements in the mouse genome. Nature 515, 355–364.

Zhang, Y., Liu, T., Meyer, C.A., Eeckhoute, J., Johnson, D.S., Bernstein, B.E., Nusbaum, C., Myers, R.M., Brown, M., Li, W., et al. (2008). Model-based analysis of ChIP-Seq (MACS). Genome Biol 9, R137.

